# Myosin II regulatory light chain phosphorylation and formin availability modulate cytokinesis upon changes in carbohydrate metabolism

**DOI:** 10.1101/2022.09.20.508671

**Authors:** Francisco Prieto-Ruiz, Elisa Gómez-Gil, Rebeca Martín-García, Armando Jesús Pérez-Díaz, Jero Vicente-Soler, Alejandro Franco, Teresa Soto, Pilar Pérez, Marisa Madrid, José Cansado

**Affiliations:** Yeast Physiology Group. Department of Genetics and Microbiology. Campus de Excelencia Internacional de Ámbito Regional (CEIR) Campus Mare Nostrum, Universidad de Murcia. 30071 Murcia, Spain.; The Francis Crick Institute, 1 Midland Road, London, NW1 1AT, UK.; Instituto de Biología Funcional y Genómica (IBFG), Consejo Superior de Investigaciones Científicas, Universidad de Salamanca. 37007 Salamanca, Spain.

**Keywords:** Cytokinesis, Respiration, Myosin II, Formin, p21-activated kinase, Fission Yeast

## Abstract

Cytokinesis, which achieves the separation of daughter cells after mitosis completion, relies in animal cells on a contractile actomyosin ring (CAR), made of actin and class II myosins, whose activity is heavily influenced by regulatory light chain (RLC) phosphorylation. However, in simple eukaryotes such as fission yeast *Schizosaccharomyces pombe*, regulation of CAR dynamics by RLC phosphorylation seems dispensable. We found that redundant phosphorylation at Ser35 of the *S. pombe* RLC homolog Rlc1 by the p21-activated kinases Pak1 and Pak2, modulates Myosin II Myo2 activity and becomes essential for cytokinesis and cell growth during respiration. Previously, we showed that the Stress Activated Protein Kinase Pathway (SAPK) MAPK Sty1 controls fission yeast CAR integrity by downregulating formin For3 levels (Gomez-Gil et al.,2020). Here we report that reduced availability of formin For3-nucleated actin filaments for the CAR is the main reason for the required control of myosin II contractile activity by RLC phosphorylation during respiration-induced oxidative stress. Hence, recovery of For3 levels with antioxidants bypasses the control of Myosin II function regulated by RLC phosphorylation to allow cytokinesis and cell proliferation during respiration. Therefore, a fine-tuned interplay between Myosin II function by Rlc1 phosphorylation and environmentally controlled actin filament availability is critical for a successful cytokinesis in response to a switch to a respiratory carbohydrate metabolism.

## Introduction

Cytokinesis, the final step in cell division, enables the physical separation of daughter cells after mitosis has been completed [1]. In non-muscle animal cells, this process is based on the formation of a contractile actomyosin ring (‘CAR’), made of actin filaments and non-muscle myosin II (NMII), which generates the mechanical force for actomyosin contractility [2, 3]. The prototype NMII motor unit is a complex assembled by two heavy chains, two light chains known as ELC (essential light chain), which provide structural integrity to the complex, and two regulatory light chains or RLC, which induce a change in NMII from a folded to an extended conformation to modulate its activity in response to phosphorylation [2]. Phosphorylation of RLC at Ser19 is critical for NMII to achieve an extended conformation, which results in the spontaneous and immediate formation of bipolar filaments with enhanced actin binding affinity and increased ATPase motor activity [2, 4, 5]. Accordingly, NMII is enzymatically inactive in the absence of RLC phosphorylation at this site [6], thus resulting in defective cytokinesis and an increase in multinucleate cells [7], which are phenotypes similar to those induced by deletion or pharmacological inhibition of NMII [8, 9]. ROCK (Rho-associated coiled-coil-containing kinase), CITK (citron kinase), and ZIPK (zipper-interacting protein kinase) are involved in NMII activation by RLC phosphorylation on Ser19 during its accumulation at the cleavage furrow, and CAR contraction, stabilization, and scission during cytokinesis [2]. RLC phosphorylation at additional sites besides Ser19, including Thr18, Ser1/2 and/or Tyr-155, provide further regulatory layers to modulate either positively or negatively the contractile activity of NMII within specific cellular conditions [2].

*Schizosaccharomyces pombe*, a Crabtree-positive fission yeast that can grow through either fermentative or respiratory metabolism, is a well-established model organism studying cytokinesis [10–12]. This simple eukaryote employs a CAR to achieve cellular division with two myosin-II heavy chains known as Myo2 and Myp2/Myo3 [13]. Myo2 function is essential for viability and cytokinesis during normal growth conditions, while Myp2 plays a subtle non-essential role for cytokinesis during unperturbed growth, and is specifically required for cell survival in response to saline stress [14–16]. Contrary to animal NMII, purified Myo2 does not form filaments at physiological saline concentrations, but instead adopts a unipolar organization, with the head domains exposed to the cytoplasm and its tails anchored into medial precursor nodes of the CAR during the mitotic onset [17–20]. The essential formin Cdc12 is responsible for the nucleation and elongation of actin filaments at the nodes. At early anaphase, a search, capture, pull, and release mechanism driven by Myo2 promotes the fusion of the equatorial nodes to form a mature CAR [21–23]. For3, a non-essential diaphanous-like formin that assembles actin cables for cellular transport, also plays a significant role in fission yeast cytokinesis nucleating actin filaments for CAR assembly and disassembly [24, 25]. Remarkably, in response to cytoskeletal damage and environmental cues, the stress-activated signaling protein kinase pathway (SAPK), and its core effector Sty1, a p38 MAPK ortholog, promotes CAR disassembly and block cell division by reducing For3 levels and the availability of actin filaments [25]. Once formed, coordinated CAR constriction by Myo2 and Myp2, and concomitant plasma membrane and primary septum formation at late anaphase, generate the physical barriers that allow the separation of the two daughter cells [10, 13, 26].

Cdc4 and Rlc1 are the respective ELC and RLC shared by Myo2 and Myp2 in fission yeast [27–29]. Early evidence indicated that Ser35 and Ser36 of Rlc1, which are homologous to RLC Thr18 and Ser19 in NMIIs, are phosphorylated by the p21/Cdc42-activated kinase (PAK) ortholog Pak1/Shk1/Orb2 [30], which has also been involved in RLC phosphorylation at Ser19 in animal cells [31]. It was described that Pak1-dependent phosphorylation of Rlc1 at Ser35 and Ser36 delays cytokinesis, and that expression of a non-phosphorylatable mutant version at both residues (*rlc1-S35A S36A*), results in premature CAR constriction [30]. These observations are consistent with *in vitro* data showing that Rlc1 phosphorylation decreases the interaction of Myo2 with actin in force generation [17]. This and a later work identified Ser35 as the sole target for Pak1 both *in vitro* and *in vivo* [17, 32]. On the other hand, another study described that Myo2 motility is reduced in fission yeast cells expressing an *rlc1-S35A S36A* mutant version, and that phosphorylation at these sites has a positive effect on CAR constriction dynamics [33]. Hence, while the essential role of RLC phosphorylation for NMII activity is firmly established in animal cells, the biological significance of Rlc1 phosphorylation at Ser35 during fission yeast cytokinesis remains currently unclear.

Here we show that modulation of Myo2 activity by Rlc1 phosphorylation at Ser35 is essential for fission yeast cytokinesis and proliferation during respiratory growth. This posttranslational modification is not only exerted by Pak1 but also by Pak2, a second PAK ortholog whose expression increases during respiration. We also show that Rlc1 phosphorylation at Ser35 becomes essential due to the reduced availability of For3-nucleated actin filaments caused by SAPK activation during respiration-induced oxidative stress. Our findings reveal how formin-dependent actin filament nucleation and Myosin II activity are linked for optimal cytokinesis control in response to changes in MAPK signaling and carbon source metabolism.

## Results

### Myosin-II regulatory light chain phosphorylation is essential for S. pombe cytokinesis and growth during respiration

To gain further insight into the contribution of RLC phosphorylation during Myosin II-dependent control of cytokinesis in *S. pombe*, we expressed a C-terminal HA-tagged version of Rlc1 under the control of its native promoter in *rlc1Δ* cells. This construct was fully functional and suppressed the defects associated with the lack of Rlc1 function, including defective CAR positioning and multiseptation (Figure 1—figure supplement 1A) [27]. We noted that in extracts from exponentially growing cells the Rlc1-HA fusion migrates in SDS-PAGE as two discernible bands (Figure 1A). Rlc1 mobility in extracts from a strain expressing a mutated version where Ser36 was changed to alanine (Rlc1(S36A)-HA) was similar to the wild-type strain. In contrast, only the faster-migrating band was observed in mutants expressing either Rlc1(S53A)-HA or Rlc1 (S35A S36A)-HA fusions (Figure 1A). Thus, in these assays the slower motility band corresponds to the *in vivo* Rlc1 isoform phosphorylated at Ser35. Increased Rlc1 expression does not alter CAR integrity and/or cytokinesis in fission yeast [34]. Hence, to precisely follow Rlc1 phosphorylation and localization dynamics during the cell cycle, we co-expressed Rlc1-GFP (genomic fusion) and Rlc1-HA (integrative fusion) tagged versions in cells carrying an analogue-sensitive version of the Cdk1 kinase ortholog Cdc2 (*cdc2-asM17*) [35], and a Pcp1-GFP fusion (pericentrin SPB component; internal control for mitotic progression). Simultaneous live fluorescence microscopy and Western blot analysis of synchronized cells released from the G2 arrest, showed that *in vivo* Rlc1 phosphorylation at Ser35 was very low at the nodes stage during CAR assembly, raised gradually during ring maturation, and reached its maximum at the onset of CAR contraction until the final stages, decreasing slowly during septum closure and cell separation (Figure 1B). As early suggested [30], these results confirm that *in vivo* Rlc1 phosphorylation at Ser35 is enhanced during CAR constriction and septum formation stages.

**Figure 1.**
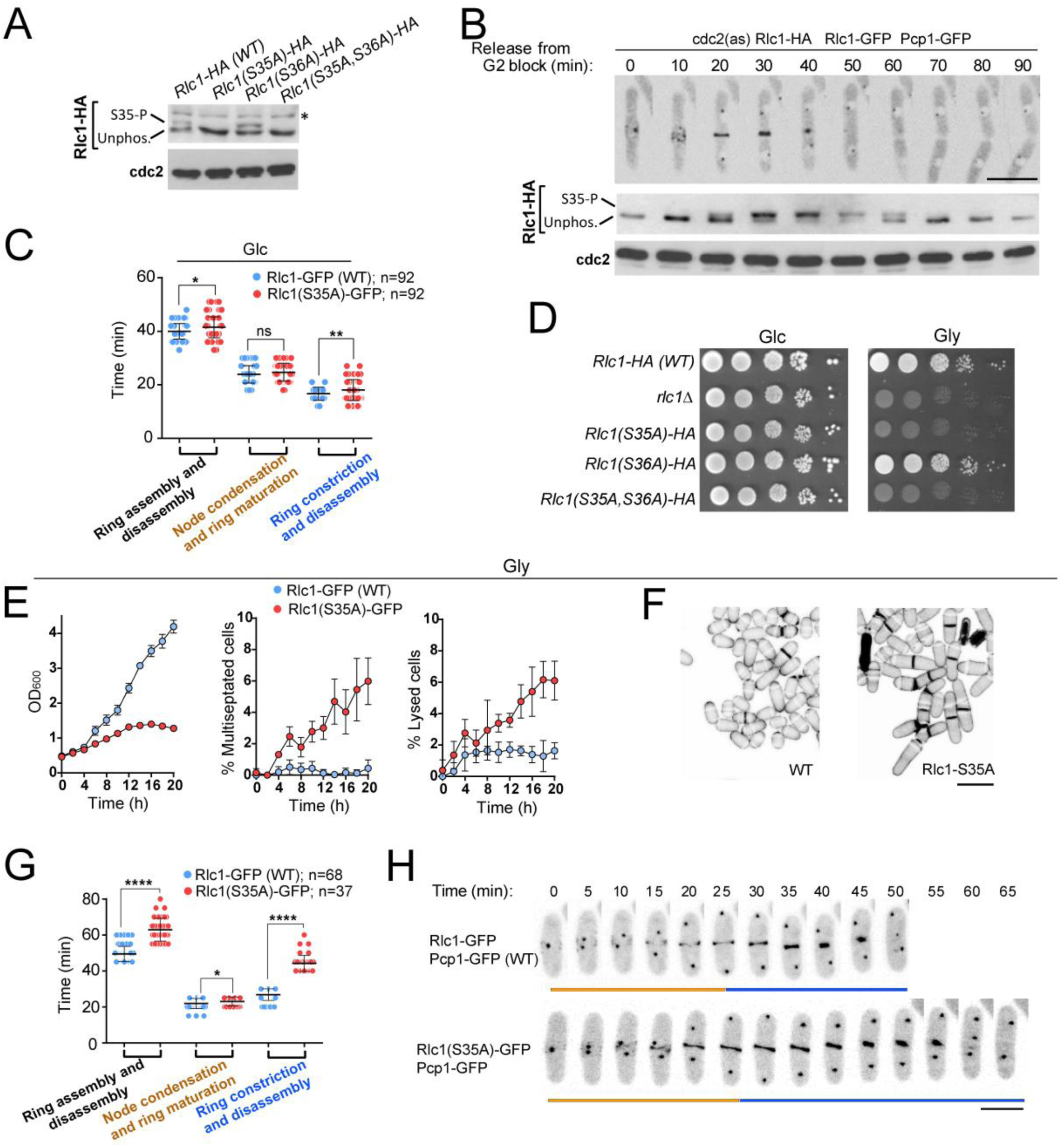
Myosin-II regulatory light chain phosphorylation is essential for *S. pombe* cytokinesis and growth during respiration. (**A**) Total protein extracts from the indicated strains growing exponentially in YES-Glucose medium were resolved by SDS-PAGE, and the Rlc1-HA fusion was detected by incubation with anti-HA antibody. Anti-Cdc2 was used as a loading control. Rlc1 isoforms, phosphorylated (S35-P), and not phosphorylated at Ser35 (Unphos), are indicated. The blot corresponds to a representative experiment that was repeated at least three times with identical results. (**B**) Cells with *cdc2-asM17* analog-sensitive mutant allele co-expressing Rlc1-HA and Rlc1-GFP genomic fusions were arrested at G2 in YES-Glucose medium supplemented with 3-NM-PP1 and incubated in the same medium lacking the kinase analogue for the indicated times. Time-lapse images of a representative cell showing Rlc1-GFP localization and mitotic progression monitored using Pcp1-GFP marked SPBs (upper panel) (scale bar: 10 µm), and Western blot analysis of Rlc1-HA mobility after release from the G2 block (lower panels), are shown. The Western blot image corresponds to a representative experiment that was repeated at least three times with identical results. (**C**) Times for ring assembly and disassembly, node condensation/ring maturation, and ring constriction and disassembly were estimated for the indicated strains growing exponentially in YES-Glucose medium (Glc), by time-lapse confocal fluorescence microscopy. *n* is the total number of cells scored from three independent experiments, and data are presented as mean ± SD. Statistical comparison between two groups was performed by unpaired Student’s *t*-test. **, p<0.005; *, p<0.05; ns, not significant, as calculated by unpaired Student’s *t* test. (**D**) Decimal dilutions of strains of the indicated genotypes were spotted on solid plates with YES-Glucose (Glc), or YES-Glycerol (Gly), incubated at 30°C for 3 (glucose plates) or 5 days (glycerol plates), and photographed. The image corresponds to a representative experiment that was repeated at least three times with similar results. (**E**) The indicated strains were grown in YES-Glucose to a final OD_600_=0.2 and shifted to YES-Glycerol and incubated at 28°C. The OD_600_ value, and the percentage of multiseptated and lysed cells were quantified in aliquots taken at the indicated times. Data correspond to three independent growth curves and are presented as mean ± SD. (**F**) Representative maximum projection confocal images of cells growing for 12 h in YES-Glycerol after cell-wall staining with calcofluor white. Scale bar: 10 µm (**G**) The times for total ring assembly and disassembly, node condensation/ring maturation, and ring constriction were estimated for the indicated strains cells growing exponentially in YES-Glycerol medium by time-lapse confocal fluorescence microscopy. Mitotic progression was monitored using Pcp1-GFP-marked SPBs. *n* is the total number of cells scored from three independent experiments, and data are presented as mean ± SD. Statistical comparison between two groups was performed by unpaired Student’s *t*-test. ****, p<0.0001; *, p<0.05, as calculated by unpaired Student’s *t*-test. (**H**) Representative maximum-projection time-lapse images of Rlc1 dynamics at the equatorial region of cells growing YES-Glycerol. Mitotic progression was monitored using Pcp1-GFP-marked SPBs. Time interval is 5 min. Scale bar: 10 µm.

Time-lapse fluorescence microscopy of exponentially glucose-growing cells from asynchronous cultures showed only a minimal but statistically significant increase in the overall time for ring constriction and disassembly in Rlc1(S35A)-GFP cells as compared to wild-type Rlc1-GFP cells (18,00 ± 3,90 *vs* 16,70 ± 2,44 min, respectively; n=92 cells) (Figure 1C). Therefore, in contrast to animal non-muscle cells, where regulatory light chain phosphorylation is essential for NMII activity [2], *in vivo* phosphorylation of Rlc1 has a minimal impact on myosin II function during CAR dynamics when fission yeast cells grow vegetatively in the presence of glucose. These findings prompted us to search for other environmental and/or nutritional condition/s where Rlc1 phoshorylation-dependent control of Myosin II activity might become essential for fission yeast cytokinesis. A recent study performed with a prototroph *S. pombe* deletion-mutant collection described that *rlc1Δ* cells struggle to grow in a glycerol-based medium that imposes a respiratory metabolism [36]. Indeed, and in contrast to wild-type cells, the growth of *rlc1Δ* cells was strongly reduced when cultured in plates with 3% glycerol and 0.08% glucose as carbon sources (Figure 1D). Strikingly, cells expressing the unphosphorylated *rlc1-S35A* and *rlc1-S35A S36A* mutants also grew very slowly in this medium (Figure 1D). This behavior was strictly dependent upon Rlc1 phosphorylation at Ser35, since expression of an Rlc1-HA fusion in *rlc1Δ* cells by employing a β-estradiol-regulated promoter [37], allowed their growth on glycerol only in the presence of 0.5 µM β-estradiol, whereas conditional expression of the unphosphorylated Rlc1(S35A)-HA mutant version did not (Figure 1—figure supplement 1B-C). Contrary to cells expressing wild-type Rlc1, the growth of *rlc1-S35A* cells transferred to a glycerol-based liquid medium was limited to 3-4 further divisions (Figure 1E), and resulted in a progressive increase in multiseptated cells with engrossed septa and lysed cells, suggesting the existence of a cytokinetic defect (Figure 1E-F). Accordingly, time-lapse fluorescence microscopy analysis revealed that the total time for ring assembly and disassembly in *rlc1-S35A* cells incubated with glycerol for 8 hours was much longer than in those expressing wild-type Rlc1 (62,97 ± 6,39 *vs* 49,49 ± 4,24 min, respectively; n≥37 cells) (Figure 1G-H). The cytokinetic delay was mainly observed during the stage of ring constriction and disassembly (44,29 ± 4,49 *vs* 26,84 ± 3,22 min, respectively) (Figure 1G-H). The above findings support that *in vivo* Rlc1phosphorylation at Ser35 plays an essential role to modulate *S. pombe* cytokinesis and cell division during respiratory growth.

### The redundant p21/Cdc42-activated kinases Pak2 and Pak1 phosphorylate Rlc1 at Ser35 together with to positively control fission yeast cytokinesis and division during respiratory growth

Earlier work has provided strong evidence that the essential fission yeast p21 (cdc42/rac)-activated protein kinase (PAK) Pak1/Shk1/Orb2, phosphorylates Rlc1 at Ser35 both *in vitro* and *in vivo* [30, 32]. In agreement with these studies, the *in vivo* phosphorylation of an Rlc1-HA fusion at Ser35 became progressively reduced in glucose-growing cells expressing the analog-sensitive (as) kinase mutant Pak1-M460A treated with the specific kinase inhibitor 3-BrB-PP1, but not in the presence of the solvent control (Figure 2—figure supplement 1A). Interestingly, light chain phosphorylation at Ser35 was absent in glucose-growing cells expressing the hypomorphic mutant allele Pak1-M460G (Figure 2A) [30], suggesting that this kinase version is constitutively inactive towards Rlc1. Unexpectedly, Rlc1 remained phosphorylated at Ser35 in cells with this mutated kinase during growth with glycerol as a carbon source (Figure 2A). The simplest explanation for these observations is that other kinase/s besides Pak1 can specifically phosphorylate Rlc1 *in vivo* at Ser35 during respiratory growth.

**Figure 2.**
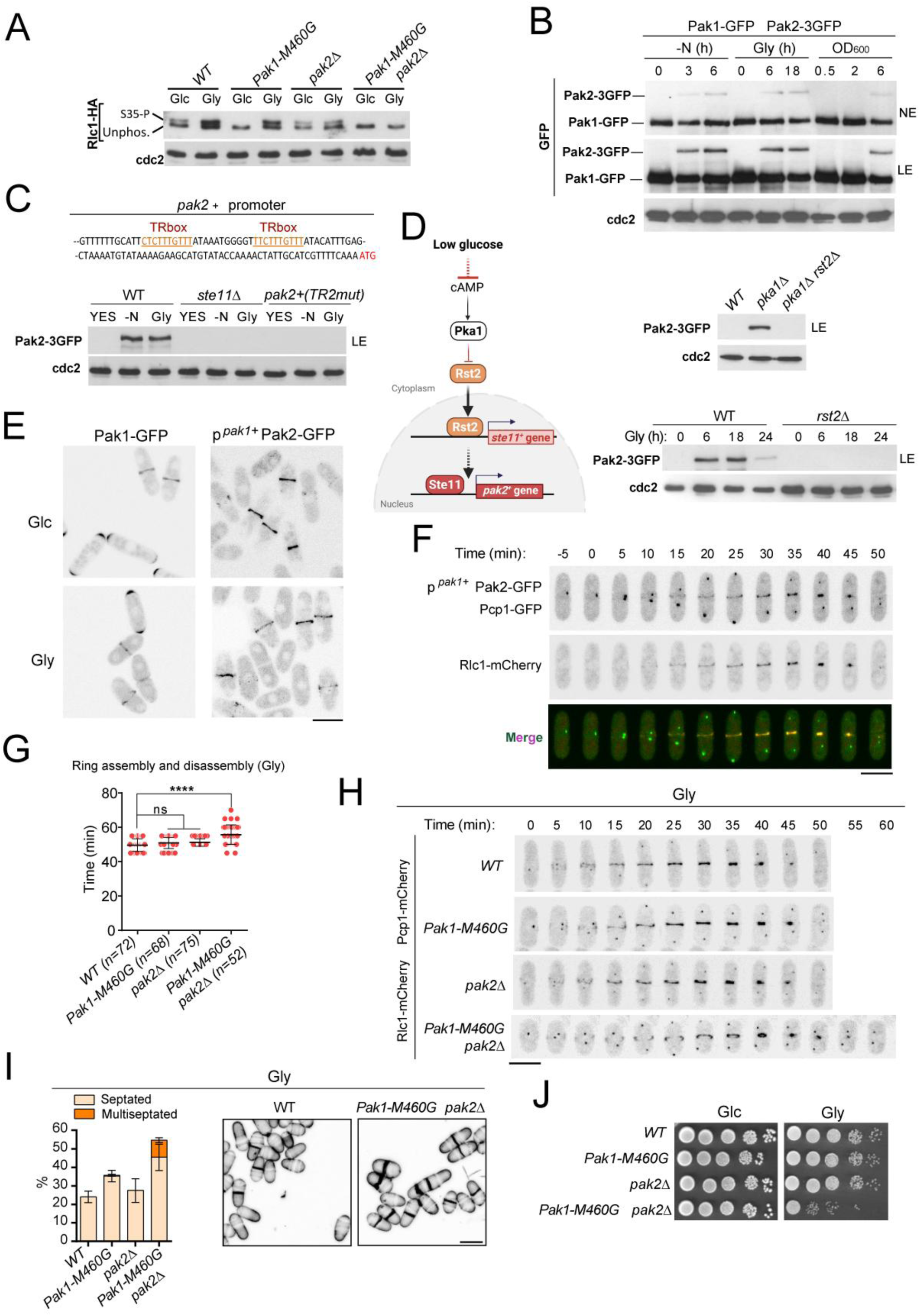
p21/Cdc42-activated kinase Pak2 phosphorylates Rlc1 at Ser35 together with Pak1 to positively control fission yeast cytokinesis and division during respiratory growth. (**A**) Total protein extracts from strains of the indicated genotypes growing exponentially in YES-Glucose (Glc) or YES-Glycerol (Gly), were resolved by SDS-PAGE, and the Rlc1-HA fusion was detected by incubation with anti-HA antibody. Anti-Cdc2 was used as a loading control. Rlc1 isoforms, phosphorylated (S35-P), and not phosphorylated at Ser35 (Unphos), are indicated. The image corresponds to a representative experiment that was repeated at least three times with similar results. (**B**) Glucose-growing cells of a *S. pombe* strain co-expressing Pak1-GFP and Pak2-3GFP genomic fusions were starved from nitrogen (-N), incubated in YES-Glycerol for the indicated times, or incubated in YES-Glucose medium until reaching the indicated OD_600_ values. The corresponding total protein extracts were resolved by SDS-PAGE, and Pak1-GFP and Pak2-3GFP fusions were detected by incubation with anti-GFP antibody. Anti-Cdc2 was used as a loading control. The image corresponds to a representative experiment that was repeated at least three times with identical results. NE: bands observed after 5 min film exposure. LE: immunoreactive bands observed after an extended film exposure of 25 min. (**C**) Upper: Partial nucleotide sequence of the promoter region of the *pak2^+^* gene. The two putative TR boxes (Ste11-binding motifs) are shown in color. Lower: Western blot analysis of Pak2-3GFP levels in wild-type, *ste11Δ*, and a mutant strain were the conserved G in the two putative TR boxes in the promoter of *pak2^+^* gene was replaced by A, growing in YES-Glucose, after nitrogen starvation (-N), and a shift to YES-Glycerol (Gly) for 12 h. Pak2-3GFP was detected by incubation with anti-GFP antibody, while anti-Cdc2 was used as a loading control. The image corresponds to a representative experiment that was repeated at least three times with identical results. (**D**) Left: Pak2 expression increases specifically during respiratory growth in a Rst2- and Ste11-dependent manner in absence of cAMP-PKA signaling. See text for a detailed description of its main components and functions. Upper right: total protein extracts from strains of the indicated genotypes growing exponentially in YES-Glucose were resolved by SDS-PAGE, and the Pak2-3GFP fusion was detected by incubation with anti-GFP antibody. Anti-Cdc2 was used as a loading control. The image corresponds to a representative experiment that was repeated at least three times with identical results. Lower right: total protein extracts from strains growing exponentially in YES-Glycerol (Gly) for the indicated times were resolved by SDS-PAGE, and the Pak2-3GFP fusion was detected by incubation with anti-GFP antibody. Anti-Cdc2 was used as a loading control. The image corresponds to a representative experiment that was repeated at least three times with identical results. (**E**) Representative maximum projection confocal images of exponentially growing cells from Pak1-GFP and p*^pak1+^*-Pak2-GFP cells growing exponentially in YES-Glucose (Glc), or YES-Glycerol (Gly). (**F**) Representative maximum-projection time-lapse images of Pak2 and Rlc1 dynamics at the CAR in cells co-expressing p*^pak1+^*-Pak2-GFP and Rlc1-mCherry genomic fusions and growing in YES-Glycerol. Mitotic progression was monitored using Pcp1-GFP-marked SPBs. Time interval is 5 min. (**G**) The total time for ring assembly and disassembly was estimated for the indicated strains growing exponentially in YES-Glycerol medium by time-lapse confocal fluorescence microscopy. Mitotic progression was monitored using Pcp1-GFP-marked SPBs. *n* is the total number of cells scored from three independent experiments, and data are presented as mean ± SD. Statistical comparison between groups was performed by one-way ANOVA. ****, p<0.0001; ns, not significant. (**H**) Representative maximum-projection time-lapse images of Rlc1 dynamics at the equatorial region in cells growing YES-Glycerol. Mitotic progression was monitored using Pcp1-GFP-marked SPBs. Time interval is 5 min. Scale bar: 10 µm. (**I**) Left: strains were grown in YES-Glycerol for 12 h, and the percentage of septated and multiseptated cells were quantified. Data correspond to three independent experiments, and are presented as mean ± SD. Right: representative maximum projection confocal images of cells from the indicated strains after cell-wall staining with calcofluor white. (**J**) Decimal dilutions of strains of the indicated genotypes were spotted on plates with YES-Glucose or YES-Glycerol, incubated at 30°C or 5 days, and photographed. The image corresponds to a representative experiment that was repeated at least three times with similar results.

A reasonable candidate to perform such a role is Pak2, a second PAK homolog whose over-expression has been shown to restore the viability and normal morphology of fission yeast cells lacking Pak1, thus suggesting that both kinases may share common substrates and functions [38, 39]. In support of this hypothesis, Rlc1 phosphorylation at Ser35 was absent during respiratory growth in Pak1-M460G *pak2Δ* double mutant cells, whereas it remained phosphorylated in a *pak2Δ* mutant growing with either glucose or glycerol (Figure 2A). In contrast to Pak1, Pak2 was undetectable after Western blot analysis in glucose-growing cells co-expressing genomic Pak1-GFP and Pak2-3GFP fusions (Figure 2B). However, its expression levels increased when the cells were either transferred to a medium lacking a nitrogen source or cultured with glycerol as a carbon source (Figure 2B). Enhanced Pak2-3GFP expression was also observed in glucose-rich medium as the cells reached the stationary phase of growth, when the availability of this carbohydrate is minimal (Figure 2B). Nevertheless, the relative expression of Pak2 was always very low compared to Pak1 and could only be detected after long exposure times of immunoblots (>20 min; LE; Figure 2B). Microarray-based studies have shown that *pak2^+^* mRNA levels increase in *S. pombe* during nitrogen starvation through a mechanism that relies on Ste11, a master transcription factor that activates gene expression during the early steps of the sexual differentiation [40]. Accordingly, *pak2Δ* cells display defective fusion during mating and produce aberrant asci [41]. The *pak2^+^* promoter shows two consecutive copies of a putative Ste11-binding motif known as the TR box (consensus sequence 5’-TTCTTTGTTY-3’) (Figure 2C) [42]. Indeed, the induced expression of a Pak2-3GFP fusion during nitrogen starvation or growth with glycerol was totally abrogated in *ste11Δ* cells, and in a strain where *pak2^+^* expression is under the control of an endogenous promoter version mutated at the two TR boxes (Figure 2C). The Zn-finger transcriptional factor Rst2, whose activity is negatively regulated by the cAMP-PKA signaling pathway in the presence of glucose, positively regulates *ste11^+^* expression during nitrogen or glucose starvation [43]. In the presence of glucose, and in contrast to wild-type cells, Pka1 deletion prompted a constitutive increase in Pak2-GFP expression (Figure 2D). Moreover, Rst2 deletion suppressed the enhanced expression of Pak2-GFP in glucose-growing *pka1Δ* cells and in the presence of glycerol (Figure 2D). Hence, Pak2 expression is constitutively repressed by cAMP-PKA signaling during glucose fermentation and increases specifically during respiratory growth in an Rst2- and Ste11-dependent manner.

The very low expression levels of the Pak2-3GFP genomic fusion prevented the microscopic visualization of its subcellular localization during nutrient starvation. To circumvent this situation, we obtained a strain expressing a Pak2-GFP fusion under the control of the native *pak1^+^* promoter (p*^pak1+^*-Pak2-GFP). The relative expression levels of p*^pak1+^*-Pak2-GFP during vegetative growth were approximately 2-3 times those of the Pak1-GFP genomic fusion (Figure 2—figure supplement 1B). Nevertheless, in contrast to Pak1-GFP, which is targeted at the cell poles and the CAR during vegetative growth with either glucose or glycerol, the p*^pak1+^*-Pak2-GFP fusion localized exclusively to the CAR in both conditions (Figure 2E). Time-lapse fluorescence microscopy of glycerol-growing cells revealed that Pak2 co-localized with Rlc1 during the entire cytokinetic process, starting with the early steps of CAR assembly and maturation to the later stages of ring constriction (Figure 2F).

Compared to wild-type cells, Pak1-M460G and *pak2Δ* cells did not display cytokinetic, septation, or growing defects during respiration with glycerol as a carbon source (Figure 2G-J). Strikingly, cells from a Pak1-M460G *pak2Δ* double mutant showed a noticeable increase in the average time for CAR assembly and disassembly (Fig 2G,H), a multiseptated phenotype (Figure 2I), and a moderated growth defect in glycerol-rich medium with respect to the wild type, Pak1-M460G, and *pak2Δ* single mutants (Figure 2J). Taken together, our observations support that Pak1 is fully responsible for *in vivo* Rlc1 phosphorylation at Ser35 during fermentative growth, whereas Pak2, whose expression is induced upon nutrient starvation, collaborates with Pak1 to phosphorylate Rlc1 at this residue to regulate cytokinesis during respiratory growth.

### PAK phosphorylation of Rlc1 becomes critical for S. pombe cytokinesis during respiration due to impaired For3-dependent actin cable nucleation imposed by SAPK activation

The formin For3 assembles actin cables for cellular transport and co-operates with the essential formin Cdc12 to nucleate actin filaments for fission yeast CAR assembly and disassembly [24, 25]. For3 absence elicits a clear delay during ring constriction and/or disassembly in glucose-rich medium [24]. The cytokinetic delay of *for3Δ* cells increased significantly in the presence of glycerol (Figure 3—figure supplement 1A), and, similar to the *rlc1-S35A* mutant (Figure 1E-F), led to the accumulation of multiseptated and lysed cells and a marked growth defect (Figure 3—figure supplement 1B-D), These observations support that actin cable nucleation by For3 is a crucial factor ensuring proper cytokinesis and growth of *S. pombe* cells during respiratory metabolism.

Glucose limitation or absence causes the activation of Sty1, a p38 MAPK ortholog, and the critical effector of the fission yeast SAPK pathway [44]. We have recently shown that activated Sty1 down-regulates CAR assembly and stability in *S. pombe* in response to cytoskeletal damage and environmental stress by reducing For3 levels [25]. Notably, the transfer of exponentially growing cells co-expressing genomic Sty1-HA and For3-3GFP fusions from a glucose-rich medium to a medium with glycerol, induced a rise in Sty1 activity that was maintained over time, as measured by Western blot analysis with anti-phospho-p38 antibody, and a concomitant decrease in For3-3GFP protein levels (Figure 3A). Similar to environmental stresses [25], the drop in For3 levels observed during growth in glycerol is likely associated with increased ubiquitination of the formin, since it was attenuated in cells of the temperature-sensitive proteasome mutant *mts3-1* (Figure 3—figure supplement 1E). We used immunofluorescence microscopy of cells stained with AlexaFluor-488-phalloidin, and calculated the number and density of actin cables by computing the cable-to-patch ratio via image segmentation using the machine learning routine Ilastik [45]. As shown in Figure 3B, the actin cable/patch ratio was significantly lower in wild-type cells growing with glycerol than in those growing with glucose as carbon source. The actin patches also appeared partially depolarized and their density increased during growth with glycerol (Figure 3B). Fluorescence intensity at the cell poles and the CAR of a CRIB-3GFP probe that detects explicitly the activated state of Cdc42 GTPase, which triggers For3 activation *in vivo* [46], was also decreased under respiratory growth conditions (Figure 3—figure supplement 2A-B). Accordingly, the overall intensity of a For3-3GFP fusion at the cell poles (G2 cells), and the CAR (late M cells), was reduced during respiratory growth as compared to glucose-fermenting cells (Figure 3B).

**Figure 3.**
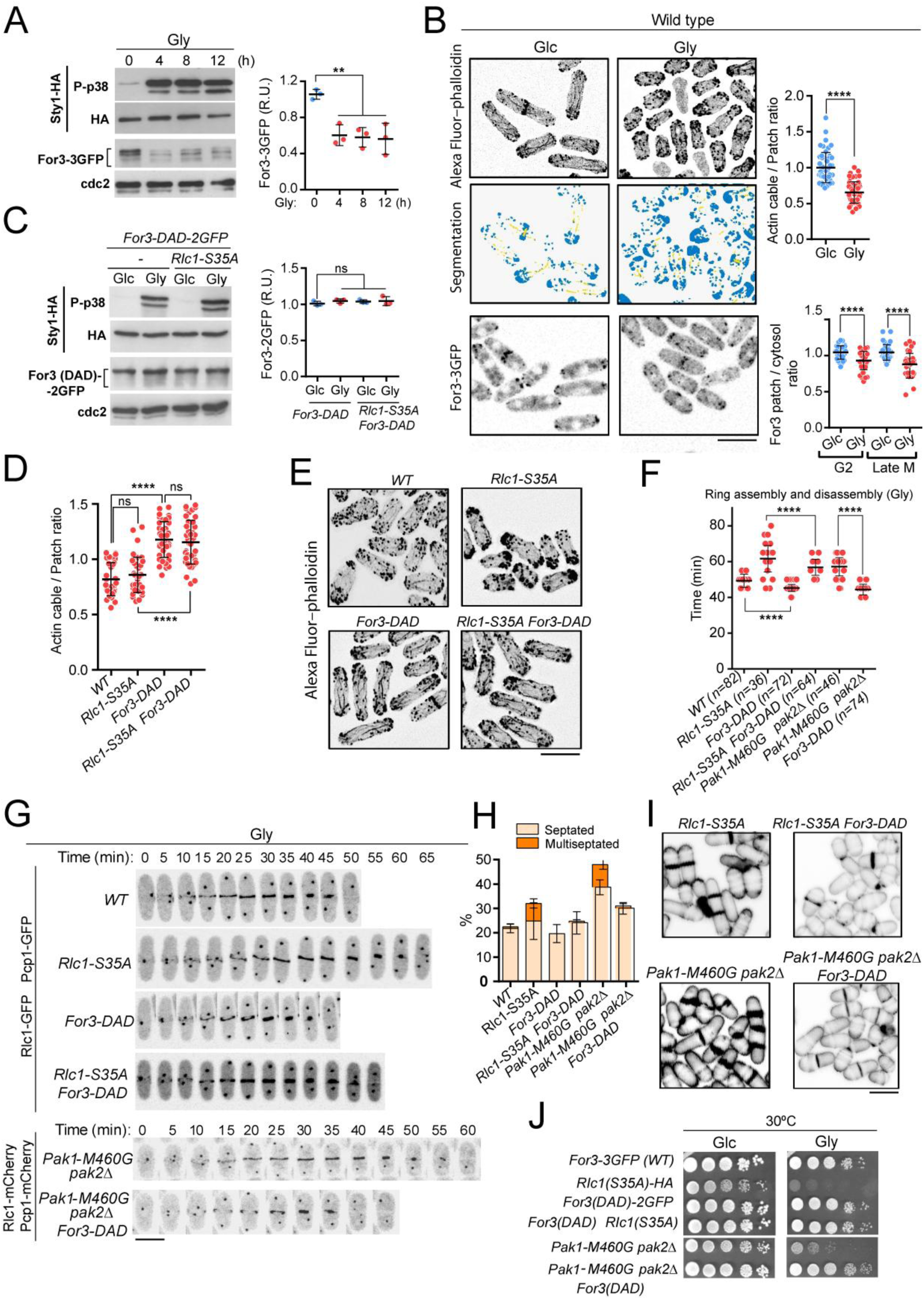
PAK phosphorylation of Rlc1 is critical for *S. pombe* cytokinesis during respiration due to impaired For3-dependent actin cable nucleation imposed by SAPK activation. (**A**) Left: *S. pombe* cells expressing genomic Sty1-HA and For3-3GFP fusions were grown in YES-Glucose to mid-log phase and transferred to YES-Glycerol medium for the indicated times. Activated/total Sty1 were detected with anti-phospho-p38 and anti-HA antibodies, respectively. Total For3 levels were detected with anti-GFP antibody. Anti-Cdc2 was used as a loading control. Right: For3 expression levels are represented as mean relative units ± SD and correspond to experiments performed as biological triplicates. **, p<0.005, as calculated by unpaired Student’s *t*-test. (**B**) Upper: representative maximum projection images of Alexa Fluor–phalloidin stained *S. pombe* cells growing in YES-Glucose medium (Glc), or in YES-Glycerol (Gly) for 12 h. Segmentation analysis with Ilastik routine is shown below each image. Quantification data correspond to the actin cable to patch ratio of G2 cells (n=51) growing with Glucose or Glycerol, and are represented as mean relative units ± SD. ****, p<0.0001, as calculated by unpaired Student’s *t*-test. Lower: representative maximum projection images of *S. pombe* cells expressing a genomic For3-3GFP fusion growing in YES-Glucose medium or in YES-Glycerol for 12 h. Quantification data (mean relative units ± SD), correspond to the For3 patch to cytosol ratio of G2 and late M cells (n=36), growing with Glucose or Glycerol. ****, p<0.0001, as calculated by unpaired Student’s *t*-test. Scale bar: 10 µm. (**C**) Left: *S. pombe* wild-type and *rlc1-S35A* strains expressing genomic Sty1-HA and For3 (DAD)-2GFP fusions were grown in YES-Glucose (Glc) to mid-log phase and transferred to YES-Glycerol medium (Gly) for 12 h. Activated/total Sty1 were detected with anti-phospho-p38 and anti-HA antibodies, respectively. Total For3 levels were detected with anti-GFP antibody. Anti-Cdc2 was used as a loading control. Right: For3 expression levels are represented as mean relative units ± SD and correspond to experiments performed as biological triplicates. ns, not significant, as calculated by unpaired Student’s *t*-test. (**D**) Actin cable to patch ratio of G2 cells from the indicated strains growing in YES-Glycerol medium for 12 h. Quantification data (n=41 cells for each strain), are represented as mean relative units ± SD. ****, p<0.0001; ns, not significant, as calculated by unpaired Student’s *t*-test. (**E**) Representative maximum projection images of Alexa Fluor–phalloidin stained *S. pombe* cells of the indicated strains growing in YES-Glycerol medium for 12 h. (**F**) The total time for ring assembly and disassembly was estimated for the indicated strains growing exponentially in YES-Glycerol medium by time-lapse confocal fluorescence microscopy. *n* is the total number of cells scored from three independent experiments, and data are presented as mean ± SD. ****, p<0.0001, as calculated by unpaired Student’s *t*-test. (**G**) Representative maximum-projection time-lapse images of Rlc1 dynamics at the equatorial region in cells from the indicated strains growing in YES-Glycerol. Mitotic progression was monitored using Pcp1-GFP-marked SPBs. Time interval is 5 min. (H) The indicated strains were grown in YES-Glycerol liquid medium for 12 h, and the percentage of septated and multiseptated cells were quantified. Data correspond to three independent experiments and are presented as mean ± SD. (**I**) Representative maximum projection confocal images of cells from the indicated strains after cell-wall staining with calcofluor white. (**J**) Decimal dilutions of strains of the indicated genotypes were spotted on plates with YES-Glucose (Glc) or YES-Glycerol (Gly), incubated at 30°C or 5 days, and photographed. The image corresponds to a representative experiment that was repeated at least three times with similar results.

In sharp contrast to wild-type cells (Figure 3A), we observed that the total levels of a constitutively active genomic version of this formin (For3(DAD)−2GFP)[46] were not reduced in response to Sty1 activation during growth with glycerol (Figure 3C). Cells expressing this Sty1-insensitive For3 allele displayed engrossed actin cables with an increased actin cable to patch ratio (Figure 3D-E), and required a shorter time for CAR assembly and disassembly as compared to the wild type (Figure 3F-G). Most importantly, the simultaneous expression of the *for3-DAD* allele in *rlc1-S35A* cells suppressed to a large extent their altered cable organization (Figure 3D-E), cytokinetic delay and multiseptated phenotype (Figure 3F-I), and defective growth in glycerol-rich medium (Figure 3J). The expression of *for3-DAD* also significantly abrogated the cytokinetic and growth defects in glycerol of a Pak1-M460G *pak2Δ* double mutant (Figure 3F-J), which lacks detectable *in vivo* phosphorylation of Rlc1 at Ser35 (Figure 2A). Together, these observations indicate that For3 and PAK-phosphorylated Rlc1 may perform a collaborative role during cytokinesis that becomes biologically significant when *S. pombe* cells grow through a respiratory metabolism.

In line with our previous observations [25], total For3-GFP levels increased in glucose-growing *wis1Δ* or *rlc1-S35A wis1Δ* strains lacking the Sty1-activating MAPKK Wis1[47] (Figure 4A), and remained significantly higher than in the Sty1-activated isogenic counterparts during growth with glycerol (Figure 4A). Similar to the *sty1Δ* mutant [25], *wis1Δ* cells growing in glycerol showed engrossed actin cables and an increase in the actin cable to patch ratio per cell with respect to the wild type (Figure 4B-C). Significantly, Wis1 deletion increased the actin cable to patch ratio in Rlc1-S35A-GFP cells (Figure 4B-C), strongly suppressed their delayed cytokinesis and multiseptation (Figure 4D-G), and restored their growth in glycerol-rich medium to a large extent (Figure 4H). Phosphorylation of Rlc1 at Ser35 has a minimal biological impact on *S. pombe* CAR dynamics during glucose fermentation (Figure 1C). Our results predict that a constitutive increase in Sty1 activity should elicit cytokinetic defects in glucose-growing *rlc1-S35A* cells. In agreement with this view, *rlc1-S35A* cells lacking the MAPK tyrosine phosphatase Pyp1, which display increased basal Sty1 activity and reduced For3 levels [25], underwent a significant delay in the time for CAR assembly and closure (Figure 4I-J), and accumulated septated cells during stationary phase (Figure 4K-L). Therefore, tight control of Rlc1 function by phosphorylation at Ser35 becomes essential for *S. pombe* cytokinesis when the levels of For3 formin are reduced by activated SAPK pathway.

**Figure 4.**
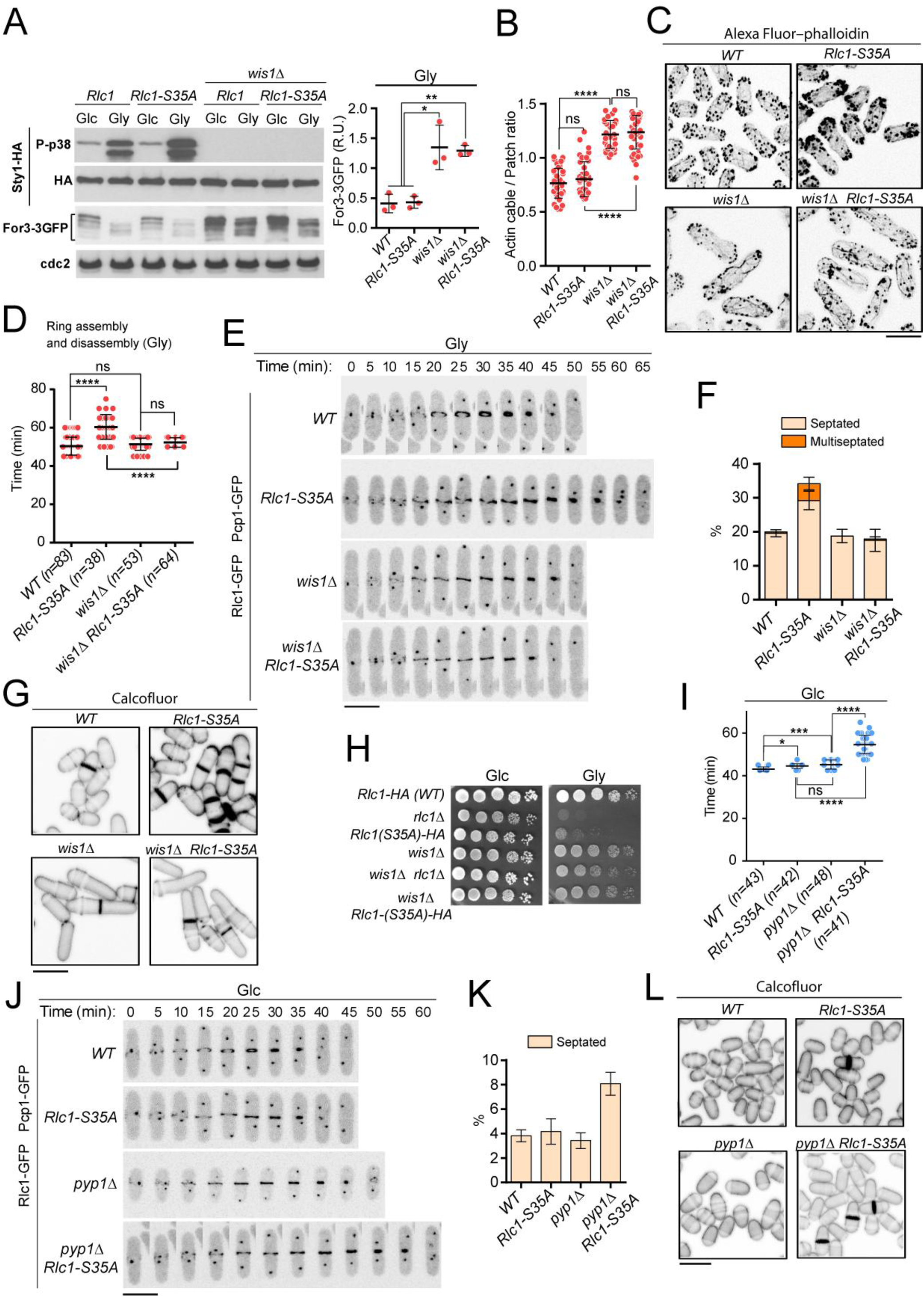
Lack of SAPK signaling restores *S. pombe* cytokinesis and growth during respiration in absence of Rlc1 phosphorylation. (**A**) Left: *S. pombe* strains of the indicated genotypes expressing genomic Sty1-HA and For3-3GFP fusions were grown in either YES-Glucose (Glc) or YES-Glycerol (Gly) medium for 12 h. Activated/total Sty1 were detected with anti-phospho-p38 and anti-HA antibodies, respectively. Total For3 levels were detected with anti-GFP antibody. Anti-Cdc2 was used as a loading control. Right: For3 expression levels in glycerol-growing strains (Gly) are represented as mean relative units ± SD and correspond to experiments performed as biological triplicates. *, p<0.05; **, p<0.005, as calculated by unpaired Student’s *t*-test. (**B**) Actin cable to patch ratio of G2 cells from the indicated strains growing in YES-Glycerol medium for 12 h. Quantification data (n=41 cells for each strain), are represented as mean relative units ± SD. ****, p<0.0001; ns, not significant, as calculated by unpaired Student’s *t*-test. (**C**) Representative maximum projection images of Alexa Fluor–phalloidin stained *S. pombe* cells of the indicated strains growing in YES-Glycerol medium for 12 h. Scale bar: 10 µm. (**D**) The total time for ring assembly and disassembly was estimated for the indicated strains growing exponentially in YES-Glycerol medium by time-lapse confocal fluorescence microscopy. *n* is the total number of cells scored from three independent experiments, and data are presented as mean ± SD. ****, p<0.0001; ns, not significant, as calculated by unpaired Student’s *t*-test. (**E**) Representative maximum-projection time-lapse images of Rlc1 dynamics at the equatorial region in cells from the indicated strains growing in YES-Glycerol. Mitotic progression was monitored using Pcp1-GFP-marked SPBs. Time interval is 5 min. (**F**) The indicated strains were grown in YES-Glycerol liquid medium for 12 h, and the percentage of septated and multiseptated cells were quantified. Data correspond to three independent experiments and are presented as mean ± SD. (**G**) Representative maximum projection confocal images of cells from the indicated strains after cell-wall staining with calcofluor white. (**H**) Decimal dilutions of strains of the indicated genotypes were spotted on plates with YES-Glucose or YES-Glycerol, incubated at 30°C or 5 days, and photographed. The image corresponds to a representative experiment that was repeated at least three times with similar results. (**I**) The total time for ring assembly and disassembly was estimated for the indicated strains growing exponentially in YES-Glucose medium by time-lapse confocal fluorescence microscopy. *n* is the total number of cells scored from three independent experiments, and data are presented as mean ± SD. ****, p<0.0001;***, p<0.001; *, p<0.05; ns, not significant, as calculated by unpaired Student’s *t*-test. (**J**) Representative maximum-projection time-lapse images of Rlc1 dynamics at the equatorial region in cells from the indicated strains growing in YES-Glucose. Mitotic progression was monitored using Pcp1-GFP-marked SPBs. Time interval is 5 min. (**K**) The indicated strains were grown in YES-Glucose liquid medium, and the percentage of septated cells were quantified. Data correspond to three independent experiments and are presented as mean ± SD. (**L**) Representative maximum projection confocal images of cells from the indicated strains after cell-wall staining with calcofluor white.

### Control of Myo2 activity by Rlc1 phosphorylation regulates S. pombe cytokinesis and growth during respiration

The thermo-sensitive and motor-deficient Myosin II heavy chain allele *myo2-E1* carries a mutation (G345R), that results in reduced ATPase activity and actin-filament binding *in vitro* [13]. We observed that the deficient growth in glycerol of *rlc1-S35A* cells was also displayed by *Myo2-E1* cells, but not by those expressing a hypomorphic allele of the essential Myosin II light chain Cdc4 (*cdc4-8*) or in a mutant lacking Myp2, a second Myosin II heavy chain that collaborates with Myo2 for CAR constriction during growth and saline stress [13, 16, 48] (Figure 5A). Accordingly, the time for CAR assembly and disassembly was much longer in *myo2-E1* cells incubated at a semi-restrictive temperature (2 h at 30°C), and changed to the permissive temperature in a glycerol-based medium as compared to glucose-rich conditions (78,59 ± 11,09 *vs* 61,67 ± 7,56 min, respectively) (Figure 5B-C). Although the respiration-induced cytokinetic delay was evident during both stages of CAR assembly/maturation and ring constriction/disassembly, it was more intense during the latter stages of ring closure (Figure 5B-C). Similar to *rlc1-S35A* cells, the *Myo2-E1* cytokinetic defect also resulted in the accumulation of many lysed and multiseptated cells with engrossed septa (Figure 5D-E). On the contrary, the delay in CAR closure displayed by glucose-growing *myp2Δ* cells *versus* wild-type cells (∼7.5 min; n≥37 cells), was very similar to that observed during growth in glycerol (∼7.7 min; n≥65 cells) (Figure 5—figure supplement 1A). Myp2 deletion increased the number of septated and multiseptated cells during growth in glucose and, particularly, in the presence of glycerol. This phenotype was aggravated when combined with the *rlc1-S35A* allele (Figure 5—figure supplement 1B).

**Figure 5.**
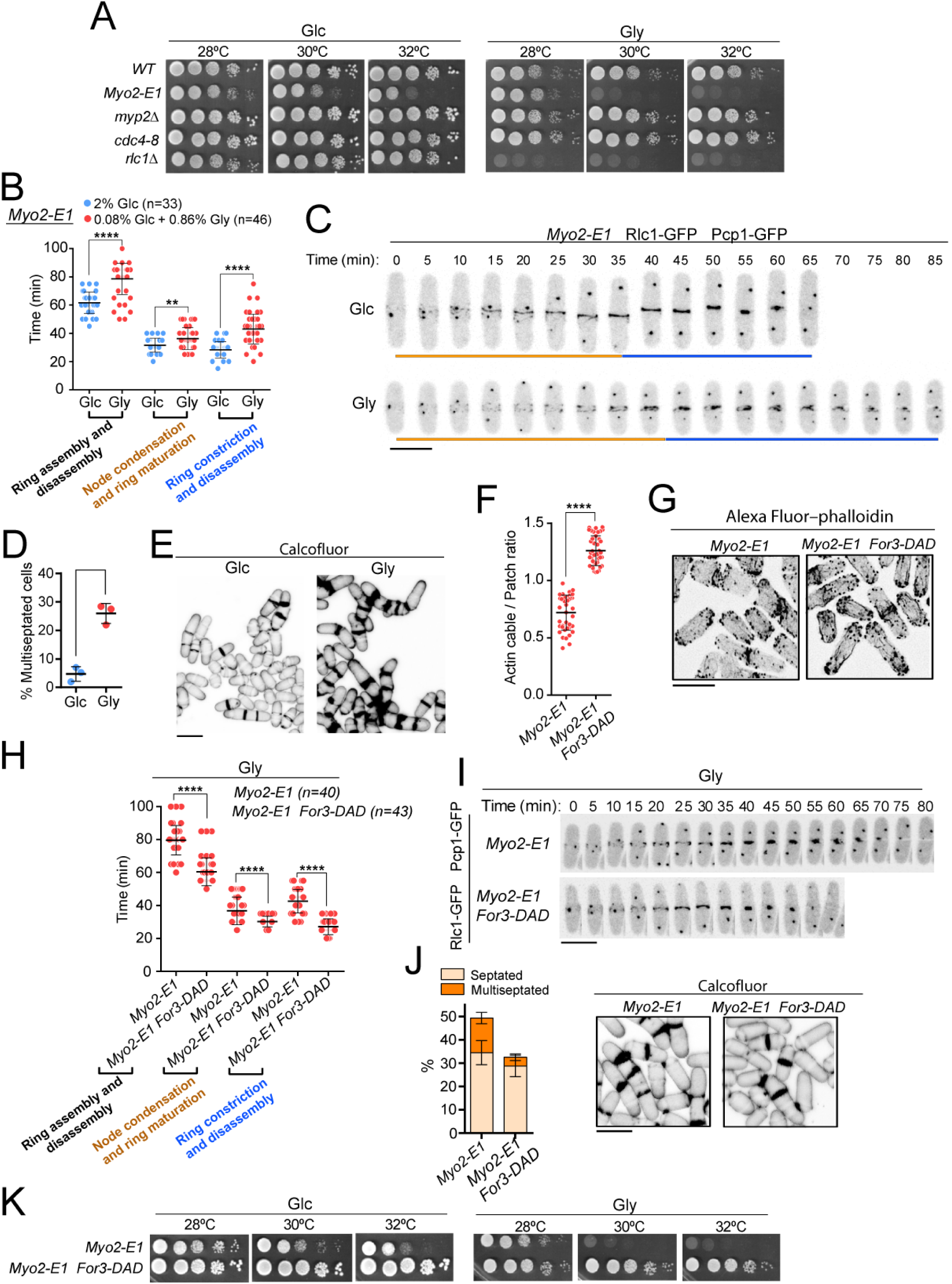
Control of Myo2 activity by Rlc1 phosphorylation regulates *S. pombe* cytokinesis and rowth during respiration. (**A**) Decimal dilutions of strains of the indicated genotypes were spotted on plates with YES-Glucose or YES-Glycerol, incubated at 28, 30, and 32°C for 3 (Glc) or 5 (Gly) days, and photographed. The images correspond to a representative experiment that was repeated at least three times with similar results. (**B**) The times for ring assembly and disassembly, node condensation/ring maturation, and ring constriction and disassembly were estimated for *Myo2-E1* cells growing in YES-Glucose (Glc) and YES-Glycerol medium (Gly), by time-lapse confocal fluorescence microscopy. Mitotic progression was monitored using Pcp1-GFP-marked SPBs. *n* is the total number of cells scored from three independent experiments, and data are presented as mean ± SD. ****, p<0.0001; *, p<0.05, as calculated by unpaired Student’s *t*-test. (**C**) Representative maximum-projection time-lapse images of Rlc1 dynamics at the equatorial region in *Myo2-E1* Rlc1-GFP cells growing in YES-Glucose (Glc) or YES-Glycerol (Gly). Mitotic progression was monitored using Pcp1-GFP-marked SPBs. Time interval is 5 min. Scale bar: 10 µm. (**D**) The percentage of multiseptated cells was quantified in *Myo2-E1* cells growing exponentially in YES-Glucose (Glc) or YES-Glycerol (Gly) for 12 h. Data correspond to three independent experiments, and are presented as mean ± SD. (**E**) Representative maximum projection confocal images of *Myo2-E1* cells growing in YES-Glucose or YES-Glycerol after cell-wall staining with calcofluor white. (**F**) Actin cable to patch ratio of G2 cells from the indicated strains growing in YES-Glycerol medium for 12 h. Quantification data (n=40 cells for each strain), are represented as mean relative units ± SD. ****, p<0.0001, as calculated by unpaired Student’s *t*-test. (**G**) Representative maximum projection images of Alexa Fluor–phalloidin stained *S. pombe* cells of the indicated strains growing in YES-Glycerol medium for 12 h. (**H**) The times for ring assembly and disassembly, node condensation/ring maturation, and ring constriction and disassembly were estimated for *Myo2-E1* and *Myo2-E1 for3-DAD* cells growing exponentially in YES-Glycerol medium by time-lapse confocal fluorescence microscopy. Mitotic progression was monitored using Pcp1-GFP-marked SPBs. *n* is the total number of cells scored from three independent experiments, and data are presented as mean ± SD. Statistical comparison between two groups was performed by unpaired Student’s *t*-test. ****, p<0.0001, as calculated by unpaired Student’s *t* test. (**I**) Representative maximum-projection time-lapse images of Rlc1-GFP dynamics at the equatorial region in *Myo2-E1* and *Myo2-E1 for3-DAD* cells growing in YES-Glycerol. Mitotic progression was monitored using Pcp1-GFP-marked SPBs. Time interval is 5 min. (**J**) Left: the percentage of septated and multiseptated cells were quantified in *Myo2-E1* and *Myo2-E1 for3-DAD* cells growing for 12 h in YES-Glycerol medium. Data correspond to three independent experiments, and are presented as mean ± SD. Right: representative maximum projection confocal images after cell-wall staining with calcofluor white. (**K**) Decimal dilutions of strains of the indicated genotypes were spotted on plates with YES-Glucose or YES-Glycerol, incubated at 28, 30, and 32°C for 3 (Glc) or 5 (Gly) days, and photographed. The images correspond to a representative experiment that was repeated at least three times with similar results.

Expression of the *for3-DAD* allele alleviated the altered cable organization (Figure 5F-G), and reduced the cytokinetic delay of *myo2-E1* cells during CAR assembly/maturation and ring constriction/disassembly (Figure 5H-I), as well as their multiseptated phenotype (Figure 5J). Moreover, simultaneous expression of *for3-DAD* restored the growth of *myo2-E1* cells in glucose- and glycerol media at semi-restrictive temperatures (Figure 5K). On the other hand, either *myo2-E1 rlc1-S35A* and *myo2-E1 wis1Δ* double mutants were synthetic lethal. Therefore, specific regulation of myosin II heavy chain Myo2 function by Pak1/Pak2-dependent *in vivo* phosphorylation of Rlc1 becomes critical to ensure *S. pombe* cytokinesis and division during respiration due to decreased For3-dependent actin filament nucleation elicited by SAPK activation.

### Exogenous antioxidants bypass the need for Rlc1 phosphorylation to regulate myosin II activity and cytokinesis during respiratory growth

In animal cells, aerobic respiration is accompanied by the production of reactive oxygen species (ROS), which typically arise due to electron leakage from the mitochondrial electron transport chain [49]. *S. pombe* cells also produce free radicals during respiratory growth [50], and the ensuing endogenous oxidative stress prompts Sty1 activation and an antioxidant response both at transcriptional and translational levels [51] (Figure 6A). Remarkably, the increased Sty1 basal activity of wild-type and *rlc1-S35A* cells growing with glycerol was largely counteracted in the presence of 0.16 mM of reduced glutathione (GSH), an antioxidant tripeptide, and this was accompanied by a recover in total For3 levels (Figure 6A). As expected, segmentation analysis confirmed that the cable-to-patch ratio was significantly improved in glycerol-growing *rlc1-S35A* and *Myo2-E1* cells supplemented with GSH, compared to those growing without the antioxidant (Figure 6B-C). Furthermore, the delayed CAR constriction and disassembly, and the multiseptation of both *rlc1-S35A* or *Myo2-E1* cells during growth with glycerol, were alleviated in the presence of GSH (Figure 6D-G). Most importantly, the simple addition of GSH allowed *rlc1-S35A* and *Myo2-E1* cells to resume growth and proliferate in the presence of glycerol (Figure 6H). This growth-recovery phenotype was For3-dependent since it was not shown by *rlc1-S35A for3Δ* cells incubated in the presence of the antioxidant (Figure 6H). Hence, oxidative stress is the leading cause of the formin-dependent reduction in the nucleation of actin cables, and imposes regulation of Myo2 function by Rlc1 phosphorylation as a critical factor in the execution of cytokinesis during respiration.

**Figure 6.**
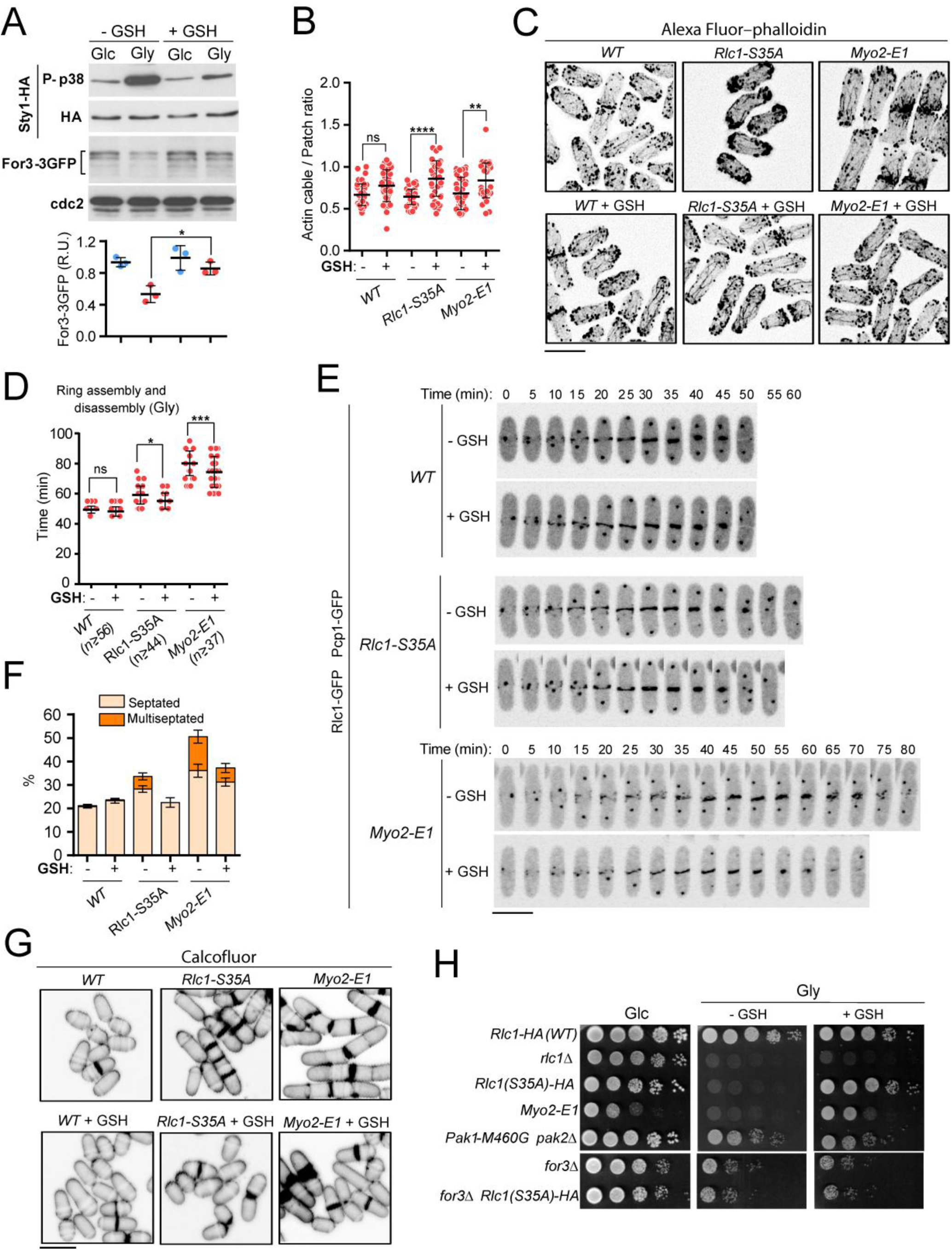
Exogenous antioxidants bypass the need for Rlc1 phosphorylation to regulate myosin II activity and cytokinesis during respiratory growth. (**A**) Upper: *S. pombe* wild type cells expressing genomic Sty1-HA and For3-3GFP fusions were grown to mid-log phase in YES-Glucose (Glc) or YES-Glycerol (Gly), with or without 0.16 mM reduced glutathione (GSH). Activated/total Sty1 were detected with anti-phospho-p38 and anti-HA antibodies, respectively. Total For3 levels were detected with anti-GFP antibody. Anti-Cdc2 was used as a loading control. Lower: For3 expression levels are represented as mean relative units ± SD and correspond to experiments performed as biological triplicates. *, p<0.05, as calculated by unpaired Student’s *t*-test. (**B**) Actin cable to patch ratio of G2 cells from the indicated strains growing in YES-Glycerol with or without 0.16 mM GSH. Quantification data (n=41 cells for each strain), are represented as mean relative units ± SD. ****, p<0.0001; **, p<0.01; ns, not significant, as calculated by unpaired Student’s *t*-test. (**C**) Representative maximum projection images of Alexa Fluor–phalloidin stained *S. pombe* cells of the indicated strains growing for 12 h in YES-Glycerol medium with or without 0.16 mM GSH. Scale bar: 10 µm. (**D**) The total time for ring assembly and disassembly was estimated for the indicated strains growing exponentially in YES-Glycerol medium with or without 0.16 mM GSH by time-lapse confocal fluorescence microscopy. *n* is the total number of cells scored from three independent experiments, and data are presented as mean ± SD. ***, p<0.001; *, p<0.05 ns, not significant, as calculated by unpaired Student’s *t* test. (**E**) Representative maximum-projection time-lapse images of Rlc1 dynamics at the equatorial region in cells from the indicated strains growing in YES-Glycerol with or without 0.16 mM GSH. Mitotic progression was monitored using Pcp1-GFP-marked SPBs. Time interval is 5 min. (**F**) The percentage of septated and multiseptated cells were quantified in the indicated strains growing for 12 h in YES-Glycerol medium with or without 0.16 mM GSH. Data correspond to three independent experiments, and are presented as mean ± SD. (**G**) Representative maximum projection confocal images of cells growing in YES-Glycerol after cell-wall staining with calcofluor white. (**H**) Decimal dilutions of strains of the indicated genotypes were spotted on plates with YES-Glucose or YES-Glycerol plates with or without 0.16 mM GSH, incubated at 28°C for 3 (Glc) or 5 (Gly) days, and photographed. The images correspond to a representative experiment that was repeated at least three times with similar results.

## Discussion

RLC phosphorylation is a common regulatory mechanism of myosin II activity both in muscle and non-muscle cells. RLC phosphorylation plays a key positive role as a regulator of myosin II function in cardiac muscle contraction under normal and disease conditions [52]. Patients with heart failure usually show reduced RLC phosphorylation, and restoring its normal phosphorylation status represents a promising approach toward improving the function of the diseased heart [53]. In non-muscle vertebrate cells, RLC phosphorylation at Ser19 is essential for NMII contractile activity during cell migration and division [2, 7]. This also applies to invertebrates like *Drosophila melanogaster*, where *in vivo* phosphorylation of *spaghetti-squash* RLC at the conserved Ser21 is critical to activate myosin II, thus avoiding embryonic lethality and severe cytokinesis defects [54]. Contrariwise, in the unicellular amoeba *Dictyostelium discoideum* RLC phosphorylation at the conserved Ser13 is not essential for regulating Myosin II function, since expression of a non-phosphorylatable S13A mutant version fully rescues the cytokinetic and developmental defects of RLC-null cells [55].

Despite also being a simple eukaryote, the effect of RLC phosphorylation on Myosin II activity during cytokinesis in the fission yeast *S. pombe* has remained elusive. While some studies indicated that lack of Rlc1 phosphorylation at the conserved Ser35 delays CAR constriction [17, 30], other works suggest just the opposite [33]. This Crabtree-positive organism uses aerobic fermentation instead of respiration for ATP production when glucose is available, whereas mitochondrial energy metabolism is significantly reduced [56]. Importantly, and to our knowledge, the experimental setup in all the published studies exploring the functional and mechanistic insights of fission yeast cytokinesis has relied on the employment of wild-type and mutant cells growing fermentatively in glucose-rich minimal or complex media. In these conditions, the impact of Rlc1 phosphorylation in cytokinesis is very modest since *rlc1-S35A* cells show only a minimal delay in CAR constriction and disassembly compared to wild-type cells. However, in this work, we show that this somehow secondary role becomes indispensable when yeast cells switch to a respiratory metabolism in the absence of glucose. In this metabolic state, lack of Rlc1 phosphorylation at Ser35 resulted in a significant delay in the dynamics of CAR assembly and disassembly, leading to a multiseptated phenotype and a decreased growth on respirable carbon sources such as glycerol. Moreover, our findings also support that Myo2, the leading Myosin II heavy chain isoform that regulates fission yeast CAR assembly and constriction under most conditions [57], is the main regulatory target for Rlc1 Ser35 phosphorylation to allow *S. pombe* cytokinesis during respiration. According to this view, the cytokinetic defects of cells expressing the hypomorphic allele *myo2-E1,* which shows reduced ATPase activity and actin-filament binding [13], become notoriously exacerbated during respiration and resemble those of *rlc1-S35A* cells. Therefore, the biological relevance of Rlc1 phosphorylation to modulate Myo2 activity *in vivo* for CAR assembly and constriction in *S. pombe* strongly depends on the carbohydrate metabolism during the transition from fermentation to respiration.

Fission yeast p21-activated kinase Pak1 phosphorylates Rlc1 at Ser35 *in vivo* in glucose-rich medium [30, 32]. However, genetic, biochemical, and cell biology evidences presented in this work support that the second PAK ortholog Pak2 collaborates with Pak1 to phosphorylate Rlc1 at this residue for adequate CAR contractility during respiratory growth (Figure 6—figure supplement 1). Accordingly, thelack of activity of both kinases, but not each one separately, resulted in cytokinetic and respiratory growth defects very similar to those shown by *rlc1-S35A* cells. Pak2 performs an important positive role during the nitrogen starvation-induced sexual development in fission yeast, and Pak2-null cells show mating defects that result in the formation of aberrant asci [41]. We found that Pak2 levels, which are not detected during fermentative growth with glucose, are markedly induced upon nitrogen starvation and during respiratory growth in the absence of glucose through a transcriptional mechanism involving the Ste11 transcription factor. Importantly, Ste11 expression is in turn activated by Rst2, another transcription factor whose activity is strongly repressed in the presence of nitrogen and glucose by the cAMP-PKA pathway [43], thus restricting Pak2 availability and function to physiological contexts where either or both nutrients are not available. Hence, it seems very likely that in such scenarios Pak2 may target multiple protein substrates, some of them in a redundant fashion with Pak1, as a booster for PAK activity. Recent phosphoproteomic screens have identified additional Pak1 substrates besides Rlc1 that function in fermentative growth during cytokinesis, including the F-BAR protein Cdc15 or the anilin-like medial ring protein Mid1, and also during polarized growth, such as the Cdc42 GEF Scd1, the RhoGAP Rga4, or the cell end marker Tea3, among others [32]. However, the observation that Pak2 only localizes to the CAR during respiratory growth, suggests that its functional redundancy with Pak1 might be restricted to cytokinesis-associated proteins.

In animal cells, both de novo actin assembly at the division site and cortical transport/flow contribute with actin filaments for the CAR [58–60]. In fission yeast cells, which lack an actin filament cortex, the CAR is assembled chiefly by Myo2 from actin filaments nucleated *de novo* at the cytokinesis nodes by the essential formin Cdc12 and partially from Cdc12-nucleating actin cables pulled from non-equatorial region [61, 62]. However, further evidence demonstrated that actin filaments nucleated by For3, the formin that assembles the actin cables that participate in polarized secretion and growth, also contribute to CAR formation in fission yeast [24, 25]. In turn, the activated SAPK pathway down-regulates in *S. pombe* CAR assembly and stability in response to stress by reducing For3 levels [25, 63]. Like animal cells, fission yeast respiratory metabolism induces endogenous oxidative stress with electron leakage from the mitochondrial electron transport chain [49].This resulted in enhanced SAPK activation, downregulation of For3 levels, and a concomitant reduction in actin filaments. Our findings strongly support that in this metabolic situation, Rlc1 phosphorylation becomes critical regulating Myo2 function during cytokinesis due to a decrease in For3-nucleated actin filaments. Accordingly, recovery of actin filaments availability during respiration by alternative strategies, including the expression of a constitutively active For3 version, limitation of For3 downregulation in SAPK-less mutants, or attenuation of the endogenous metabolic oxidative stress with antioxidants (GSH), was sufficient to restore CAR assembly/constriction and cytokinesis both in *rlc1-S35A* and *myo2-E1* mutants during respiration, thus allowing cell growth. Compared to wild-type cells, the number of actin filaments at the CAR is reduced approximately by half in *myo2-E1* cells [64], which might explain why enhanced nucleation of actin filaments by For3 alleviates their defective actin-binding and motor activity during cytokinesis.

Accumulating evidence suggests that metabolic reprogramming fuels the actin cytoskeletal rearrangements that occur during the response of cells to external forces, epithelial-to-mesenchymal transition, and cell migration. They are accompanied by glycolysis and oxidative phosphorylation alterations that provide the required energy for these rearrangements [65–67]. However, a yet unanswered question is how changes in cell metabolism prompt actin cytoskeletal remodeling [67]. Our observations reveal a sophisticated adaptive interplay between modulation of Myosin II function by Rlc1phosphorylation and environmentally controlled formin availability, which becomes critical for a successful cytokinesis during a respiratory carbohydrate metabolism (Figure 6—figure supplement 1). Altogether, these findings provide a remarkable example of how carbohydrate metabolism dictates the relative importance of different sources of actin filaments for CAR dynamics during cellular division.

## Acknowledgments

We thank Pedro M. Coll for providing yeast strains and plasmids. This research was funded by the Agencia Estatal de Investigación and Ministerio de Ciencia e Innovación, Spain, grant numbers PID2020-112569GB-I00 and PGC2018-098924-B-I00, and the Regional Government of Castile and Leon, Spain, grant number CSI150P20. European Regional Development Fund (ERDF), co-funding from the European Union. F.P.-R. and A.P.-D. are, respectively, Formación de Profesorado Universitario PhD fellows from the Ministerio de Educación y Formación Profesional and the Universidad de Murcia, Spain.

## MATERIALS AND METHODS

### Strain construction

*Schizosaccharomyces pombe* strains used in this work are listed in Supplementary Table S1. Several deletion strains were obtained from the Bioneer mutant library [68], whereas null mutants in *rlc1^+^, pka1*^+^ and *ste11*^+^ genes were obtained by ORF deletion and replacement with G418 (kanR), nourseothricin (NAT), or hygromycin B cassettes by employing a PCR-mediated strategy [69, 70], and the oligonucleotides described in Supplementary Table S2. Strains expressing different genomic fusions were constructed either by transformation or after random spore analysis of appropriate crosses in sporulation agar (SPA) medium [71]. To generate a strain expressing an integrated Rlc1-HA fusion, the *rlc1^+^* ORF plus its endogenous promoter were amplified by PCR using genomic DNA from *S. pombe* 972h^-^ wild-type strain as the template and the 5’ and 3’oligonucleotides PromRlc1(XhoI)-FWD and Rlc1-GFP(SacII)-REV (Supplementary Table S2), which include, respectively, a *Xho*I restriction site and an extended DNA sequence encoding a HA C-terminal tag plus a *Sac*II site. The *Xho*I-*Sac*II digested PCR fragment was cloned into plasmid pJK210 [72], sequenced, linearized with *Bmg*BI, and transformed into an *rlc1Δ ura4.294* strain. To obtain a strain expressing an integrative Rlc1-HA fusion under the control of the β-estradiol promoter [37], the *rlc1^+^* ORF fused to a 3’ HA tag was amplified by PCR 5’ and 3’oligonucleotides Rlc1 (SmaI)-FWD and Rlc1-HA (SacII)-REV, which include, respectively, *Sma*I and *Sac*II restriction sites. The amplified PCR product was cloned into a modified plasmid pJK210 containing a β-estradiol regulated promoter Z_3_EV [37], and the resulting construct was linearized with *Stu*I, and transformed into an *rlc1Δ ura4.294* strain. To obtain a strain expressing an integrative Rlc1-GFP fusion, DNA encoding an Rlc1-GFP fusion under the endogenous promoter was amplified by PCR using as the template a genomic DNA from a *S. pombe* strain expressing a genomic Rlc1-GFP fusion (Supplementary Table S1), and the 5’ and 3’oligonucleotides PromRlc1(XhoI)-FWD and Rlc1-GFP(SacII)-REV, which include *Sma*I and *Sac*II restriction sites, respectively. The resulting DNA fragment was cloned into plasmid pJK210, linearized with *Stu*I, and transformed into an *rlc1Δ ura4.294* strain. In all cases Ura4^+^ transformants were obtained, and the correct integration and expression of the Rlc1-HA and Rlc1-GFP fusions fusion under either the endogenous or the β-estradiol regulated promoters were verified by both PCR and Western blot analysis, respectively. To generate strains expressing Rlc1-HA and Rlc1-GFP versions with mutations at Ser35 and/or Ser36 residues to Alanine, the pJK210 plasmids described above containing either Rlc1-GFP or Rlc1-HA fusions were subjected as templates to site-directed mutagenesis by PCR, by employing specific mutagenic oligonucleotides described in Supplementary Table S2. Then mutagenized plasmids were linearized with *Bmg*BI and transformed into an *rlc1Δ ura4.294* strain.

The *S. pombe* strain expressing a genomic Pak2-3GFP fusion was obtained in two successive steps. First, the *pak2^+^* ORF plus its endogenous promoter were amplified by PCR using genomic DNA from *S. pombe* 972h^-^ wild-type strain as the template, and the 5’ and 3’oligonucleotides PromPak2(XhoI)-FWD (*Xho*I site) and Pak2GFP(SmaI/XmaI)-REV (*Sma*I site) (Supplementary Table S2). The PCR product was cloned in frame into a pJK210 plasmid containing a GFP C-terminal Tag. In a second step, this construct was linearized with *Sma*I and two additional GFP tags were added by a Gibson assembly approach. Finally, the resulting plasmid was linearized with *Bmg*BI and transformed in a *pak2Δ ura4.294* strain. To introduce the mutations at the two putative Ste11-binding motifs in Pak2 promoter (TR box), the pJK210-Pak2-3GFP plasmid was subjected to sequential site-directed mutagenesis by PCR. In this way, the conserved G in each motif was replaced by A by employing the mutagenic oligonucleotides described in Supplementary Table S2. To generate a strain producing a Pak2-GFP fusion under the control of Pak1 promoter, Pak1 5’UTR sequence was amplified by PCR using genomic DNA from *S. pombe* 972h^-^ wild-type strain and assembled by Gibson cloning to a PCR-amplified Pak2-GFP fragment and the pJK210 plasmid linearized with *Sma*I. The resulting plasmid was digested with *Bmg*BI and transformed in a *pak2Δ ura4.294* strain.

### Media and growth conditions

In experiments performed with liquid cultures, fission yeast strains were grown overnight with shaking at 28°C in YES-Glucose medium, which includes 0.6% yeast extract, 2% glucose, and is supplemented with adenine, leucine, histidine, or uracil (100 mg/liter) [73]. The next day, cultures were diluted to an OD_600_ of 0.01 and incubated until reaching a final OD_600_ of 0.2. Then, cells were recovered by filtration, washed three times, and shifted to either YES-Glucose or YES-Glycerol (0.6% yeast extract, 0.08% glucose, 0.86% glycerol, plus supplements), and incubated at 28°C for 4 h before imaging. In experiments performed with the *Myo2-E1* mutant, cells recovered from cultures at 28°C were resuspended in YES-Glucose or YES-Glycerol, incubated at 30°C for 2 h, and then at 28°C for the remainder of the experiment. In experiments with cells expressing the analogue-sensitive Cdc2 (CDK) kinase version *cdc2-asM17* [35], cells from log-phase liquid cultures in YES-Glucose (OD_600_ 0.5), were treated with 1 µM 3-NM-PP1 (Sigma-Aldrich, 529581) for 3.5 h, recovered by filtration, washed, and resuspended in YES-Glucose medium. In experiments with strains expressing an analog-sensitive Pak1 kinase version Pak1-M460A, log-phase liquid cultures were divided in two and incubated for different times in YES-Glucose medium treated with 10 µM 3-BrB-PP1 (Abcam, ab143756), or in medium lacking the analog kinase inhibitor. In nitrogen starvation experiments, strains growing exponentially in Edinburgh Minimal Medium (EMM2)[74] with 2% glucose (OD_600_ 0.5), were recovered by filtration and resuspended in the same medium lacking ammonium chloride for the indicated times. In the plate assays of stress sensitivity for growth, *S. pombe* wild-type and mutant strains were grown in YES-Glucose liquid medium to an OD_600_ of 1.2, recovered by centrifugation, resuspended in YES to a density of 10^7^ cells/ml, and appropriate decimal dilutions were spotted on YES-Glucose (2% glucose), or YES-Glycerol (0.08% glucose plus 3% glycerol), solid plates (2% agar). Plates were incubated for 3 days (YES-Glucose), or 5 days (YES-Glycerol), at different temperatures (28°C, 30°C, 32°C, and/or 34°C), depending on the experiment, and then photographed. All the assays were repeated at least three times with similar results. Representative experiments are shown in the corresponding Figures. When required, solid and or liquid media were supplemented with varying amounts of β-estradiol (Sigma-Aldrich, RPN2106), or reduced glutathione (GSH; Sigma-Aldrich, G6013).

### Microscopy analysis

For *time-lapse* imaging of CAR dynamics, 300 µl of cells growing exponentially for 4 h in YES-Glucose or YES-Glycerol liquid medium, and prepared as described above, were placed in a well from a μ-Slide eight well (Ibidi, 80826), previously coated with 10 μl of 1 mg/ml soybean lectin (Sigma-Aldrich, L2650) [25]. When required, GSH was incorporated into the medium at a final concentration of 0.3 mM. Cells were left to sediment in the culture media and attach to the well bottom for 1 min, and images were captured every 2.5 min for 2 h in YES-Glucose cultures, or every 5 min for 8 h in YES-Glycerol cultures. Experiments were performed at 28°C, and single middle planes from a set of six stacks (0.61 µm each) were taken at the indicated time points. Time-lapse images were acquired using a Leica Stellaris 8 confocal microscope with a 63X/1.40 Plan Apo objective and controlled by the LAS X software. The time for node condensation and ring maturation includes the time from SPB separation until the start of CR constriction. The time for ring constriction and disassembly includes the time from the first frame of ring constriction until the last frame where it becomes completely constricted and disassembled. The total time for ring assembly and disassembly is the sum of these two values. *n* is the total number of cells scored from at least three independent experiments. Statistical comparison between two groups was performed by unpaired Student’s *t-test*.

To perform actin staining with Alexa-Fluor phalloidin, 5 ml mid-log cultures in YES-Glucose (OD_600_ 0.5), or YES-Glycerol (OD_600_ 0.2), were grown for 12 h after media shift. Cells were fixed by shacking for 1 h with 3.7% formaldehyde in PEM buffer (10 mM EGTA; 1 mM MgSO_4_; 100 mM PIPES pH 6.9, 75 mM sucrose and 0.1% Triton X-100). After three washes with PEM, the cell pellets were resuspended in 20 µl of cold 40% methanol solution, stained with 8 µl of 5 mg/ml Alexa fluor 488-conjugated phalloidin (Thermo Fisher Scientific, A12379), and incubated in a rotary platform overnight at 4°C in the dark. Images of stained cells were acquired from samples spotted on glass slides with a Leica Stellaris 8 confocal microscope using a 100X/1.40 Plan Apo objective (7 stacks of 0.3 µm each). For actin segmentation analysis, the Ilastik routine with the Pixel classification tool [45], was trained with two representative images, one from cell growing in YES-Glucose medium, and one with cells growing in YES-Glycerol. The training involves drawing cables, patches and background in three different colors. Once the program was trained, the remaining images from the different experiments were uploaded to Ilastik to perform the segmentation routine. The resulting images were then exported to ImageJ [75], and segmented cells at G2 were analysed using the color histogram tool, obtaining the specific areas corresponding to cables and patches. The data from n ≥ 40 cells growing in YES-Glycerol were obtained for each cell by dividing the cable area by the patch area, and the ratio was normalized with respect to the average obtained from wild-type cells growing with YES-Glucose medium. To perform For3-GFP quantification Ilastik was trained drawing For3-GFP dots, GFP background and image background in three different colors. The For3-GFP patch to cytosol ratio was calculated by dividing the For3-GFP color area between the GFP background area from at least n ≥ 40 cells in G2 or late M (dividing cells) and normalized with the average of the glucose ratio.

For cell wall staining, fission yeast cells were cultured in YES-Glucose or YES-Glycerol for different times in the absence or presence of 0.3 mM of GSH. Cells from 1 ml aliquots were recovered by centrifugation, stained with 1 µl of 0.5 mg/ml calcofluor white, and images were acquired from samples spotted on glass slides with a Leica Stellaris 8 confocal microscope using a 63X/1.40 Plan Apo objective (6 stacks of 0.61 µm each). The percentage of septated (one septa), multiseptated (two or more septa), and lysed cells, was calculated at the indicated time points for each strain and condition from three independent experiments. n≥ 100 cells were counted from several images captured during each replicate.

### Western blot analysis

To detect levels of Rlc1-HA fusion and/or its phosphorylation status, fission yeast cultures were grown in YES-Glucose or YES-Glycerol as described above, and 10 ml samples were collected and precipitated with TCA [76]. Protein extracts were resolved in 15% SDS-PAGE gels, transferred to nitrocellulose blotting membranes, and immunoblotted with a mouse monoclonal anti-HA antibody (clone 12CA5; Roche, 11 583 816 001, RRID:AB_514505). Rabbit monoclonal anti-PSTAIR (anti-Cdc2; Sigma-Aldrich, 06-923, RRID:AB_310302) was used for loading control. Immunoreactive bands were revealed, respectively, with anti-mouse (Abcam, ab205719, RRID:AB_2755049), and anti-rabbit HRP-conjugated secondary antibodies (Abcam, ab205718, RRID:AB_2819160), and the ECL system (GE-Healthcare, RPN2106). For detection of Pak1-GFP and Pak2-3GFP fusions, the TCA-precipitated protein extracts were resolved in 6% SDS-PAGE gels, transferred to nitrocellulose membranes, and incubated with a mouse monoclonal anti-GFP antibody (Roche, 11 814 460 001, RRID:AB_390913), and anti-cdc2 (PSTAIR), as a loading control. To determine For3-3GFP and For3(DAD)-2GFP levels, total protein extracts from exponentially growing cultures were obtained under native conditions with lysis buffer (20 mM Tris-HCl pH 8.0, 2 mM EDTA, 100 mM NaCl, and 0.5% NP-40, plus a protease inhibitor cocktail). Proteins were resolved in 6% SDS-PAGE gels and transferred to Hybond-ECL membranes. For3-GFP fusions were detected with a mouse monoclonal anti-GFP antibody (Roche), with anti-cdc2 (PSTAIR) as a loading control. In all cases the immunoreactive bands were revealed with anti-mouse or anti-rabbit HRP-conjugated secondary antibodies and the ECL system.

To detect Sty1 phosphorylation and total protein levels in strains expressing a genomic Sty1-HA fusion, cell samples of 5 ml were collected at the indicated times and immediately centrifuged for 20 s at 3200 rpm/4°C. The cell pellets were resuspended in 1 ml of ice-cold buffer (10 mM NaPO_4_, 0.5 mM EDTA pH 7.5), transferred to 1.5 ml tubes, centrifuged at 13000 rpm/4°C, and stored at 80°C until further processing. Cell lysis was achieved in a FastPrep instrument after mixing the cell pellets with pre-chilled 0.5 mm glass beads to −20°C with ice-cold lysis buffer (20 mM Tris-HCl pH 8.0, 2 mM EDTA, 100 mM NaCl, and 0.5% NP-40 and containing a protease inhibitor cocktail) (Sigma Aldrich, P8340). The cell lysates were clarified by centrifugation at 13000 rpm/4°C for 5 min, and the protein extracts were resolved in 12% SDS-PAGE gels and transferred to nitrocellulose membranes. Dual phosphorylation of Sty1 was detected employing a rabbit polyclonal anti-phospho-p38 antibody (Cell Signaling, 9211, RRID:AB_331641). Total Sty1 was detected in *S. pombe* extracts with mouse monoclonal anti-HA antibody (12CA5, Roche). Immunoreactive bands were revealed with anti-mouse or anti-rabbit HRP-conjugated secondary antibodies (Abcam), and the ECL system.

Densitometric quantification of Western blot experiments as of 16-bit. jpg digital images of blots was performed using ImageJ [75]. The desired bands plus background were drawn as rectangles and a profile plot (peak) was obtained for each band. To reduce the background noise in the bands, each peak floating above the baseline of the corresponding peak was manually closed off using the straight-line tool. Measurement of the closed peaks was performed with the wand tool. Relative Units (R.U.) of For3 levels were estimated by determining the signal ratio of the correspondent anti-GFP (total For3) blot with respect to the anti-cdc2 blot (internal control) at each time point. Quantification data correspond to experiments performed as biological triplicates. Mean relative units ± SD are shown.

### Statistical analysis

Statistical analysis was performed using prism 6 software (Graph pad), and results are represented as mean ± SD, unless otherwise indicated. Comparisons for two groups were calculated using unpaired two-tailed Student’s t-tests, whereas comparisons of more than two groups were calculated using one-way ANOVA with Bonferroni’s multiple comparison tests. We observed normal distribution and no difference in variance between groups in individual comparisons. Statistical significance: * p<0.05; ** p < 0.005; *** p < 0.0005; **** p < 0.0001. Further details on statistical analysis are included in the figure legends.

## Data availability

All data generated or analyzed during this study are included in the manuscript and supporting files.

## Supplemental Information

**Figure 1—figure supplement 1.**
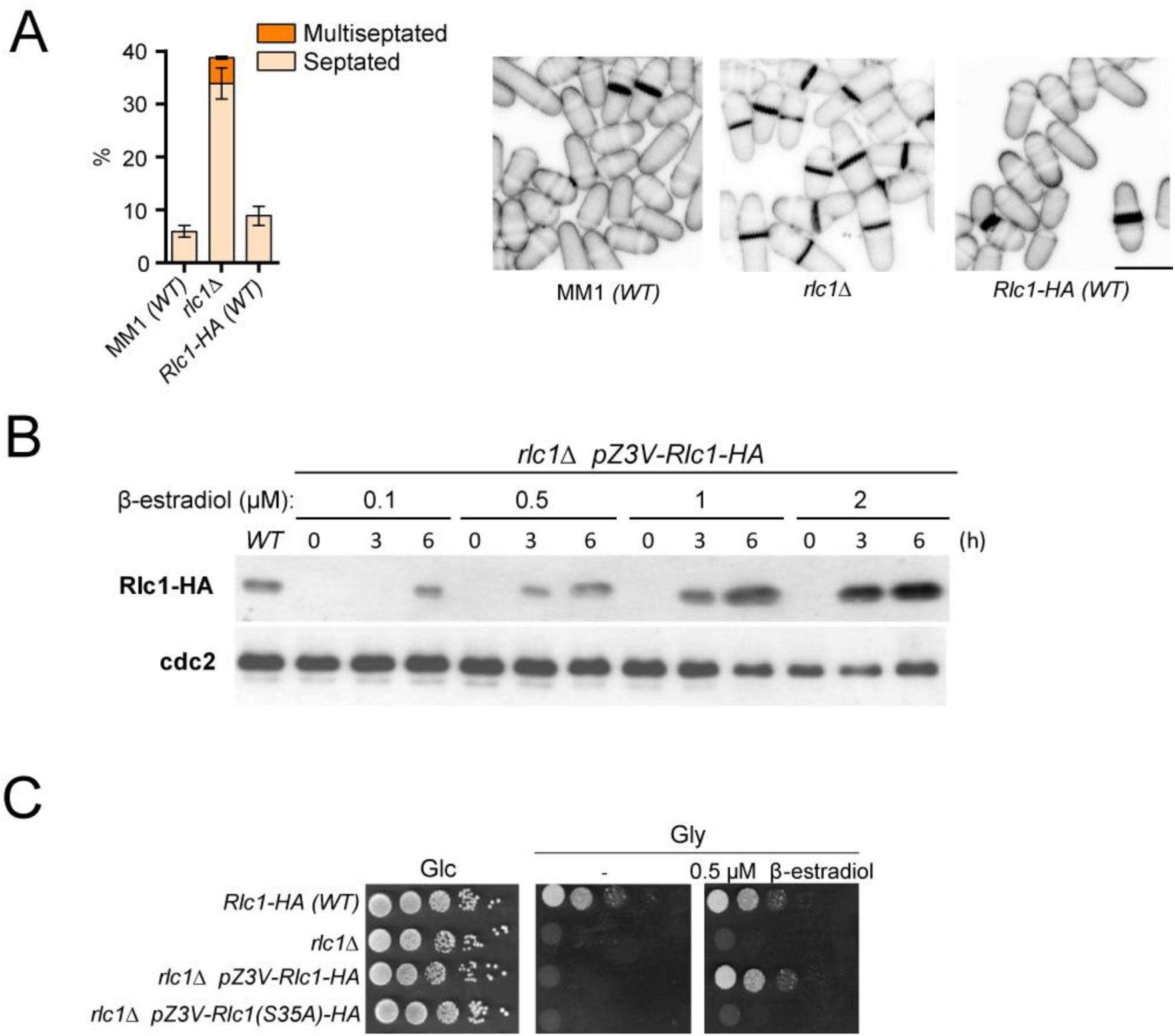
Rlc1 phosphorylation at Ser35 is essential for *S. pombe* respiratory growth. (A) Left: Strains of the indicated genotypes were grown in YES-Glucose liquid medium for 24 h, and the percentage of septated and multiseptated cells were quantified. Data correspond to three independent experiments, and are presented as mean ± SD. Right: representative maximum projection confocal images of cells from the indicated strains after cell-wall staining with calcofluor white. Scale bar: 10 µm. (B) The strain *rlc1*Δ Z3EVpr:Rlc1-HA was grown in YES-Glucose medium to mid-log phase, and the culture was then treated with either 0.1, 0.5. 1, or 2 µM β-estradiol for 0, 3 and 6 h. Total extracts were resolved by SDS-PAGE, and Rlc1 levels were detected by incubation with anti-HA antibody. Anti-Cdc2 was used as a loading control. The Western blot image corresponds to a representative experiment that was repeated at least three times with similar results. (C) Decimal dilutions of strains of the indicated genotypes were spotted on plates with YES-Glucose (Glc) or YES-Glycerol (Gly), with or without 0.5 µM β-estradiol, incubated at 30°C or 3 days, and photographed. The image corresponds to a representative experiment that was repeated at least three times with similar results.

**Figure 2—figure supplement 1.**
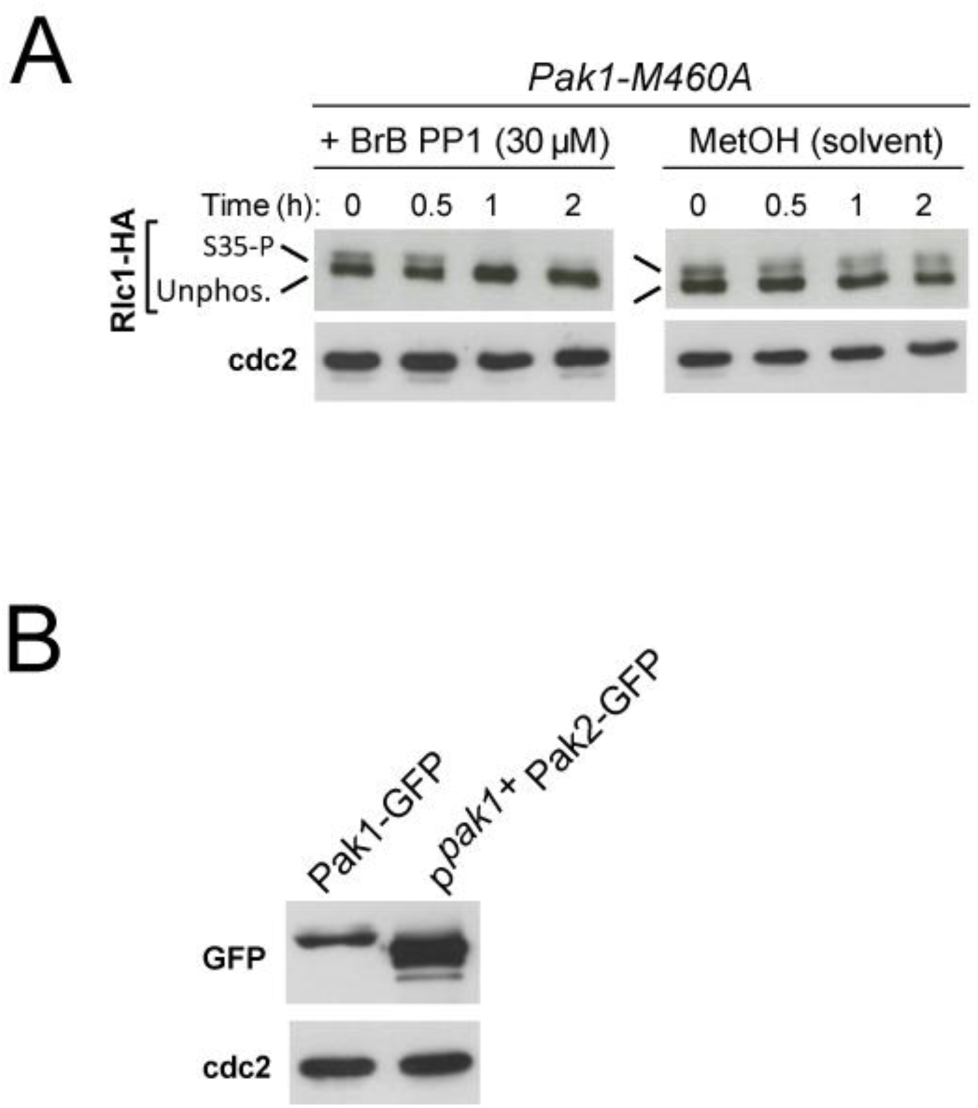
Pak1 phosphorylation of Rlc1 at Ser35 *in vivo*. (A) Exponentially growing cells of the analog-sensitive strain Pak1-M460A were grown in YES-Glucose medium and treated with 30 µM of 3-BrB-PP1 for the indicated times, or remained untreated in the presence of solvent alone (MetOH). The corresponding protein extracts were resolved by SDS-PAGE, and the Rlc1-HA fusion was detected by incubation with anti-HA antibody. Anti-Cdc2 was used as a loading control. S35-P: Rlc1 isoform phosphorylated *in vivo* at Ser35. Unphos.: Rlc1 isoform not phosphorylated at Ser35. The image corresponds to a representative experiment that was repeated at least three times with similar results. (B) Total protein extracts from strains expressing either Pak1-GFP or p*^pak1+^*-Pak2-GFP fusions and growing exponentially in YES-Glucose were resolved by SDS-PAGE. Fusions were detected by incubation with anti-GFP antibody. Anti-Cdc2 was used as a loading control. The image corresponds to a representative experiment that was repeated at least three times with similar results.

**Figure 3—figure supplement 1.**
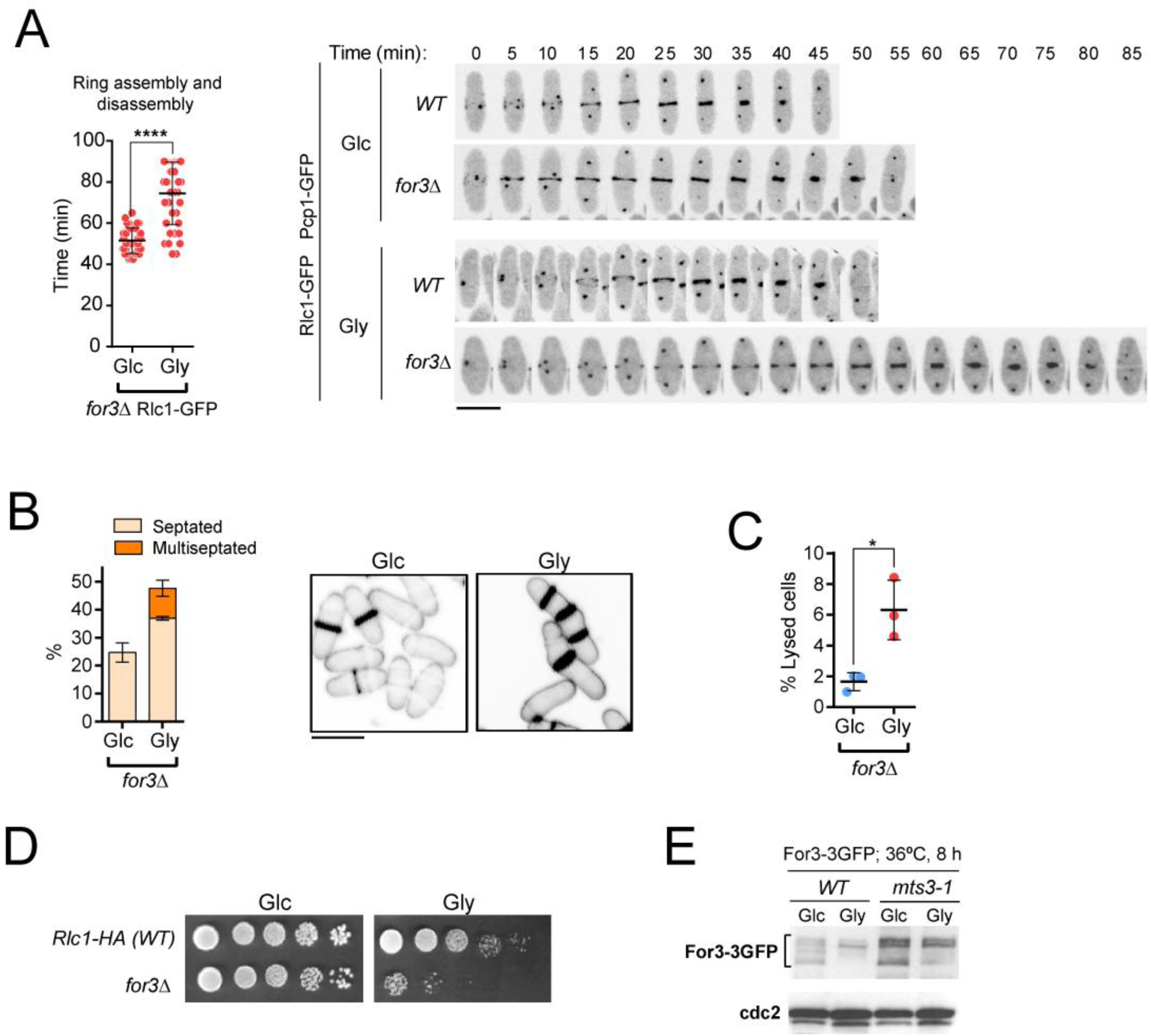
For3 formin is required for *S. pombe* cytokinesis during respiratory growth. (A) Left: the total time for ring assembly and disassembly was estimated for *for3Δ* Rlc1-GFP cells growing exponentially in either YES-Glucose (Glc) or YES-Glycerol (Gly) medium by time-lapse confocal fluorescence microscopy. n≥36 cells from three independent experiments were scored in each condition, and data are presented as mean ± SD. ****, p<0.0001, as calculated by unpaired Student’s *t* test. Right: representative maximum-projection time-lapse images of Rlc1 dynamics at the equatorial region in cells from the indicated strains growing in YES-Glucose or YES-Glycerol. Mitotic progression was monitored using Pcp1-GFP-marked SPBs. Time interval is 5 min. Scale bar: 10 µm. (B) Left: the percentage of septated and multiseptated cells were quantified in *for3Δ* cells growing for 12 h in either YES-Glucose (Glc) or YES-Glycerol medium (Gly). Data correspond to three independent experiments, and are presented as mean ± SD. Right: representative maximum projection confocal images after cell-wall staining with calcofluor white. Scale bar: 10 µm. (C) The percentage of cell lysis was quantified in *for3Δ* cells growing for 12 h in either YES-Glucose (Glc) or YES-Glycerol medium (Gly). Data correspond to three independent experiments, and are presented as mean ± SD. *, p<0.05, as calculated by unpaired Student’s *t* test. (D) Decimal dilutions of strains of the indicated genotypes were spotted on plates with YES-Glucose (Glc) or YES-Glycerol (Gly), incubated at 30°C for 5 days, and photographed. The image corresponds to a representative experiment that was repeated at least three times with similar results. (E) For3-3GFP levels were determined by Western blot analysis with anti-GFP antibody in extracts from wild-type and the proteasome mutant *mts3-1* growing in YES-Glucose (Glc) or YES-Glycerol (Gly) and incubated at 36°C for 8 h. Anti-Cdc2 was used as a loading control. The image corresponds to a representative experiment that was repeated at least three times with similar results.

**Figure 3—figure supplement 2.**
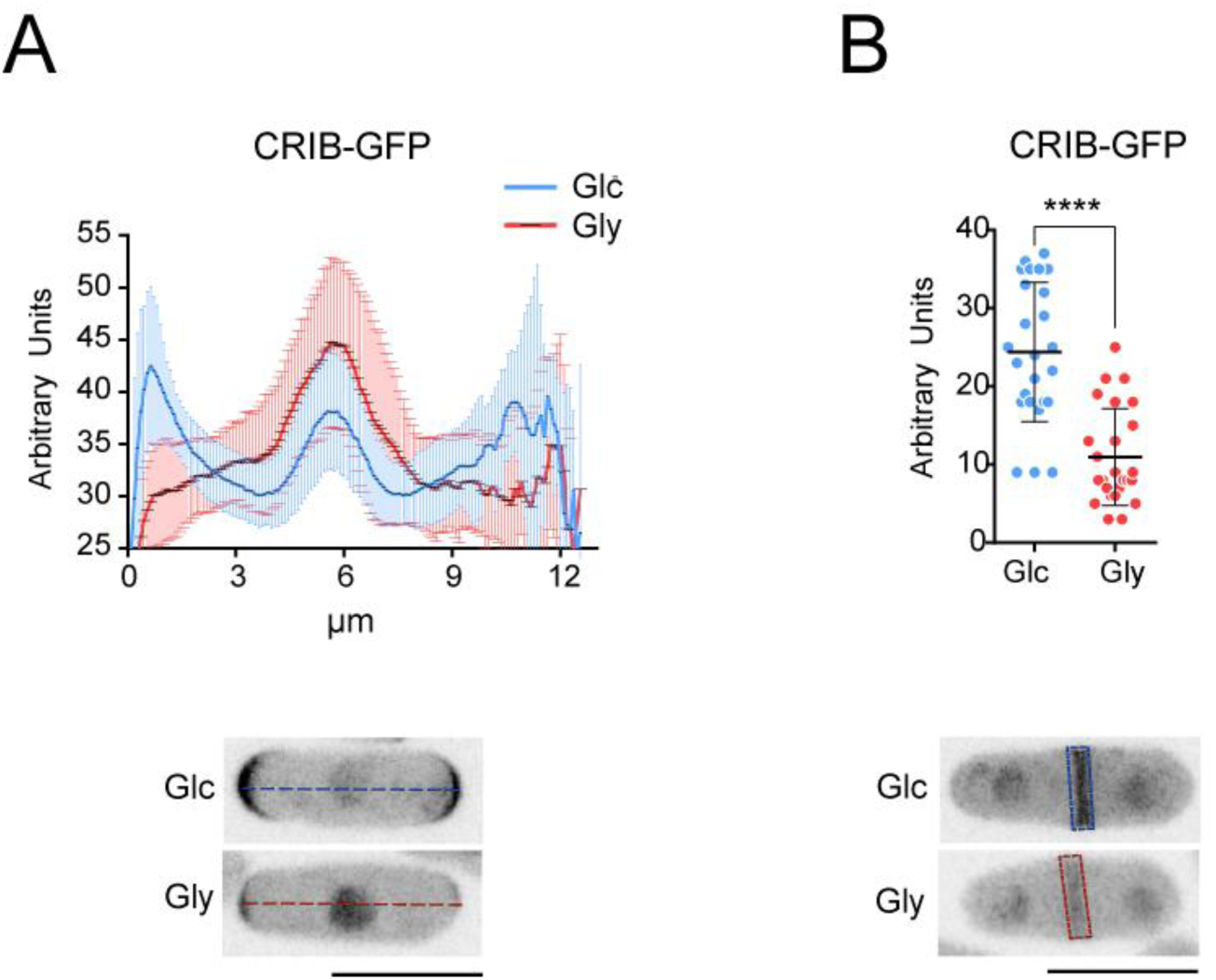
Localization of activated Ccd42 at the cell poles and the CAR is reduced during respiratory growth. (A) Upper: intensity plots of CRIB-3GFP fusion (shown as arbitrary fluorescence units), were generated from line scans across the equatorial region of *S. pombe* G2 cells (n= 30) growing in YES-Glucose (Glc) or YES-Glycerol (Gly). Data are presented as mean ± SD. Lower: representative maximum-projection images of glucose and glycerol growing cells are shown. Scale bar: 10 µm. (B) Upper: the intensity of the CRIB-3GFP fusion at the medial region of dividing cells (n= 25) in YES-Glucose (Glc) or YES-Glycerol (Gly) was measured and is shown as arbitrary fluorescence units. Data are presented as mean ± SD. ****, p<0.0001, as calculated by unpaired Student’s *t* test. Lower: representative maximum-projection images of dividing cells are shown. Scale bar: 10 µm.

**Figure 5—figure supplement 1.**
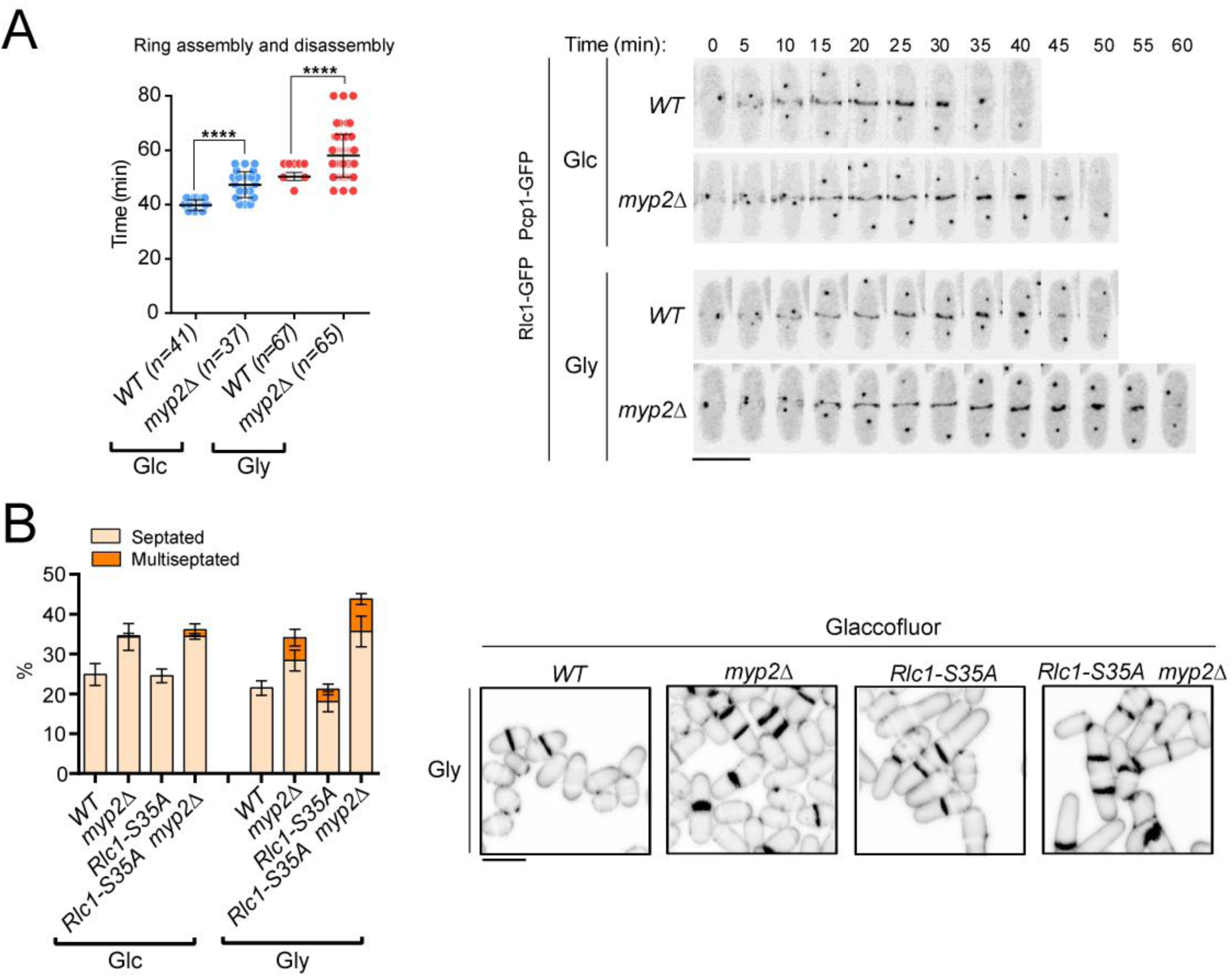
Role of Myp2 on *S. pombe* cytokinesis during respiration. (A) Left: the times for ring assembly and disassembly, was estimated for wild type and *myp2Δ* cells growing in YES-Glucose (Glc) or YES-Glycerol medium (Gly), by time-lapse confocal fluorescence microscopy. Mitotic progression was monitored using Pcp1-GFP-marked SPBs. *n* is the total number of cells scored from three independent experiments, and data are presented as mean ± SD. Statistical comparison between two groups was performed by unpaired Student’s *t* test. ****, p<0.0001, as calculated by unpaired Student’s *t* test. Right: representative maximum-projection time-lapse images of Rlc1 dynamics at the equatorial region of wild type and *myp2Δ* cells growing in YES-Glucose (Glc) or YES-Glycerol (Gly). Mitotic progression was monitored using Pcp1-GFP-marked SPBs. Time interval is 5 min. Scale bar: 10 µm. (B) Left: the percentage of septated and multiseptated cells were quantified in the indicated strains growing for 12 h in YES-Glucose (Glc) or YES-Glycerol (Gly). Data correspond to three independent experiments, and are presented as mean ± SD. Right: representative maximum projection confocal images of cells growing in YES-Glycerol after cell-wall staining with calcofluor white. Scale bar: 10 µm.

**Figure 6—figure supplement 1.**
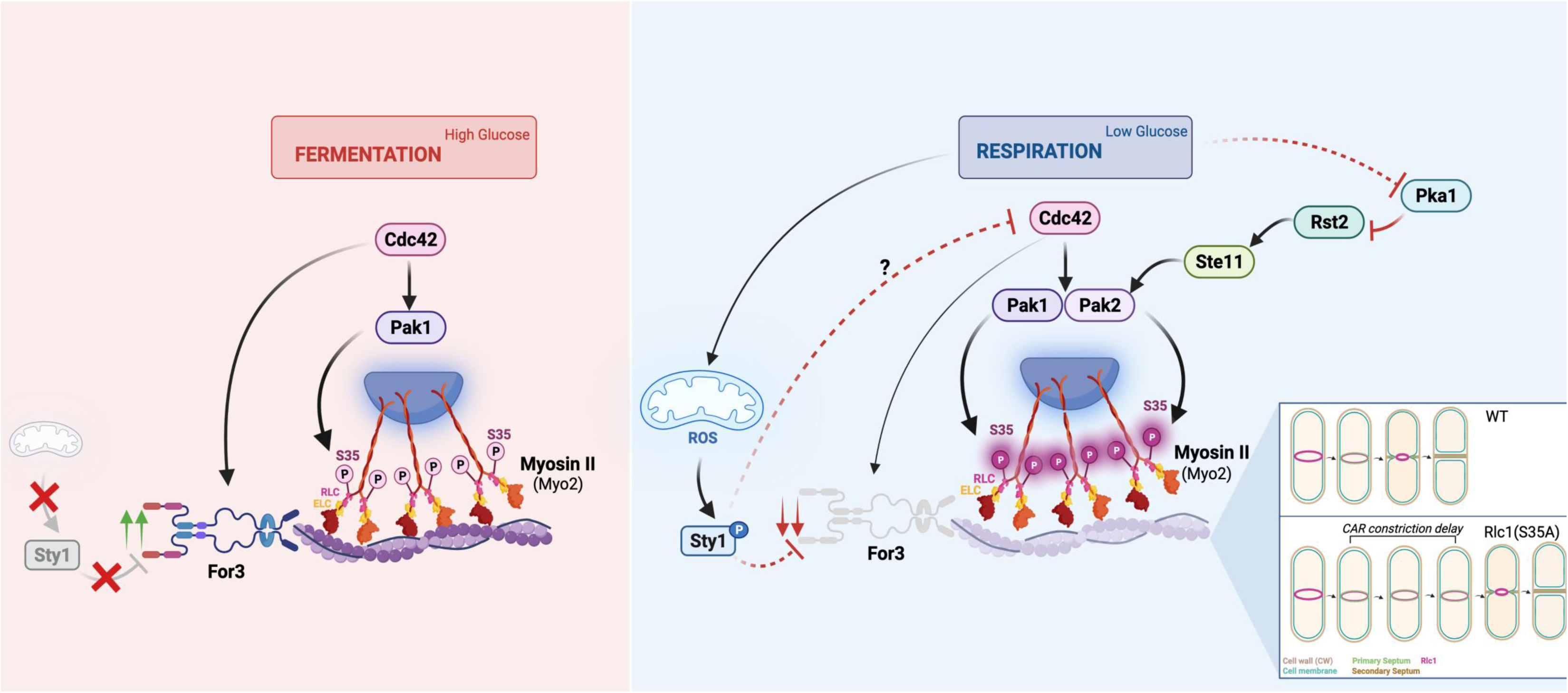
Model depicting the signaling pathways and mechanisms that regulate *S. pombe* cytokinesis by Myosin II
(Myo2) through regulatory light chain phosphorylation during the transition from fermentative to respiratory metabolism. For specific details, please see text.

## Source files legends

**Figure 1- source data 1.** Source data for Figure 1

**Figure 1- source data 2.** Western blot images for Figure 1 A,B

**Figure 1- figure supplement 1-source data 1.** Source data for Figure 1-figure supplement 1

**Figure 1- figure supplement 1-source data 2**. Western blot images for figure supplement 1B

**Figure 2- source data 1.** Source data for Figure 2

**Figure 2- source data 2.** Western blot images for Figure 2B,C,D.

**Figure 2- figure supplement 1-source data 1.** Western blot images for figure supplement 1A,B

**Figure 3- source data 1.** Source data for Figure 3

**Figure 3- source data 2.** Western blot images for Figure 3A,C.

**Figure 3- figure supplement 1-source data 1.** Source data for Figure 3-figure supplement 1

**Figure 3- figure supplement 1-source data 2.** Western blot images for figure supplement 1E

**Figure 3- figure supplement 2-source data 1.** Source data for Figure 3-figure supplement 2

**Figure 4- source data 1.** Source data for Figure 4

**Figure 4- source data 2.** Western blot images for Figure 4A.

**Figure 5- source data 1.** Source data for Figure 5

**Figure 5- figure supplement 1-source data 1.** Source data for Figure 5-figure supplement 1

**Figure 6- source data 1.** Source data for Figure 6

**Figure 6- source data 2.** Western blot images for Figure 6A.

---

**Table S1.**
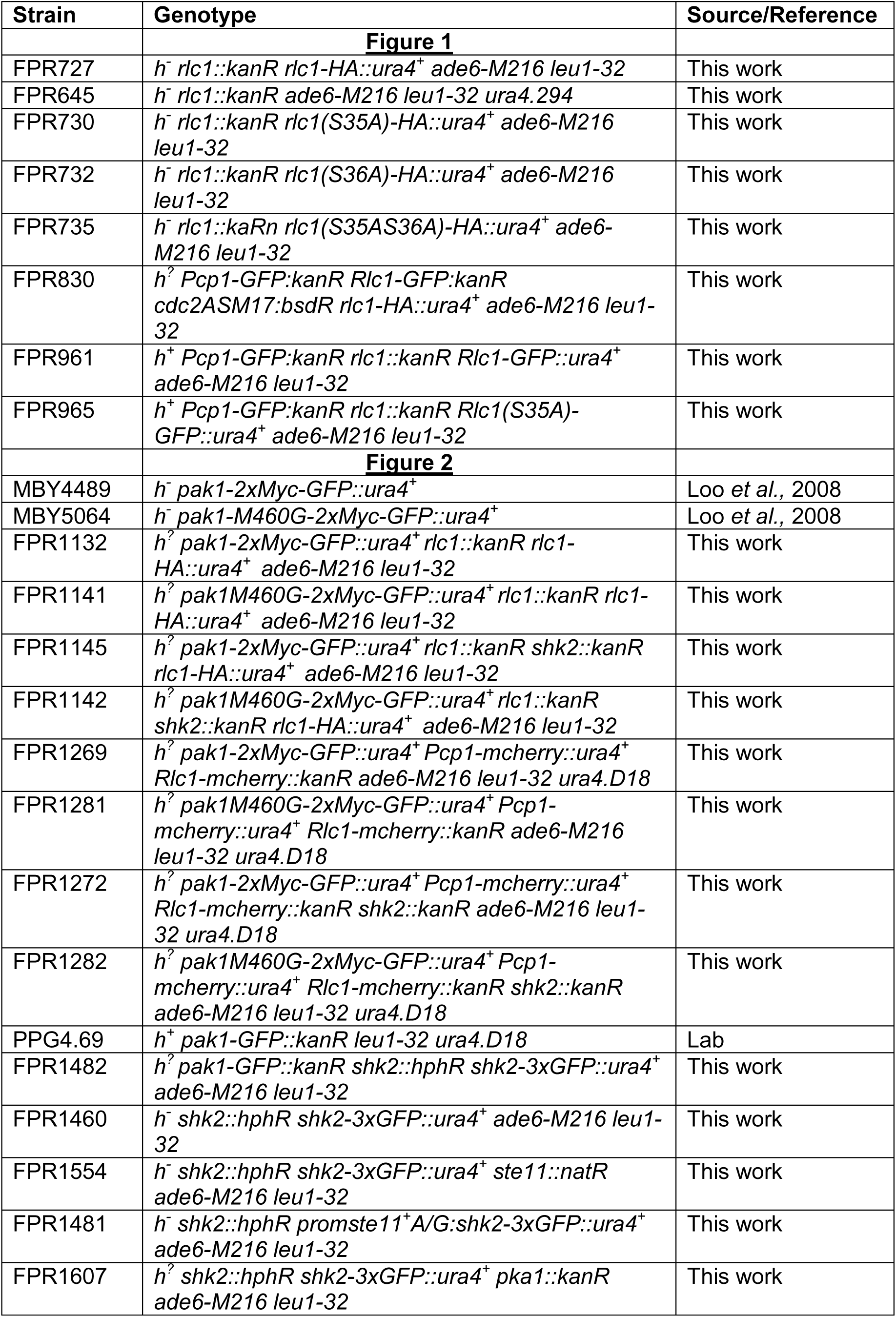
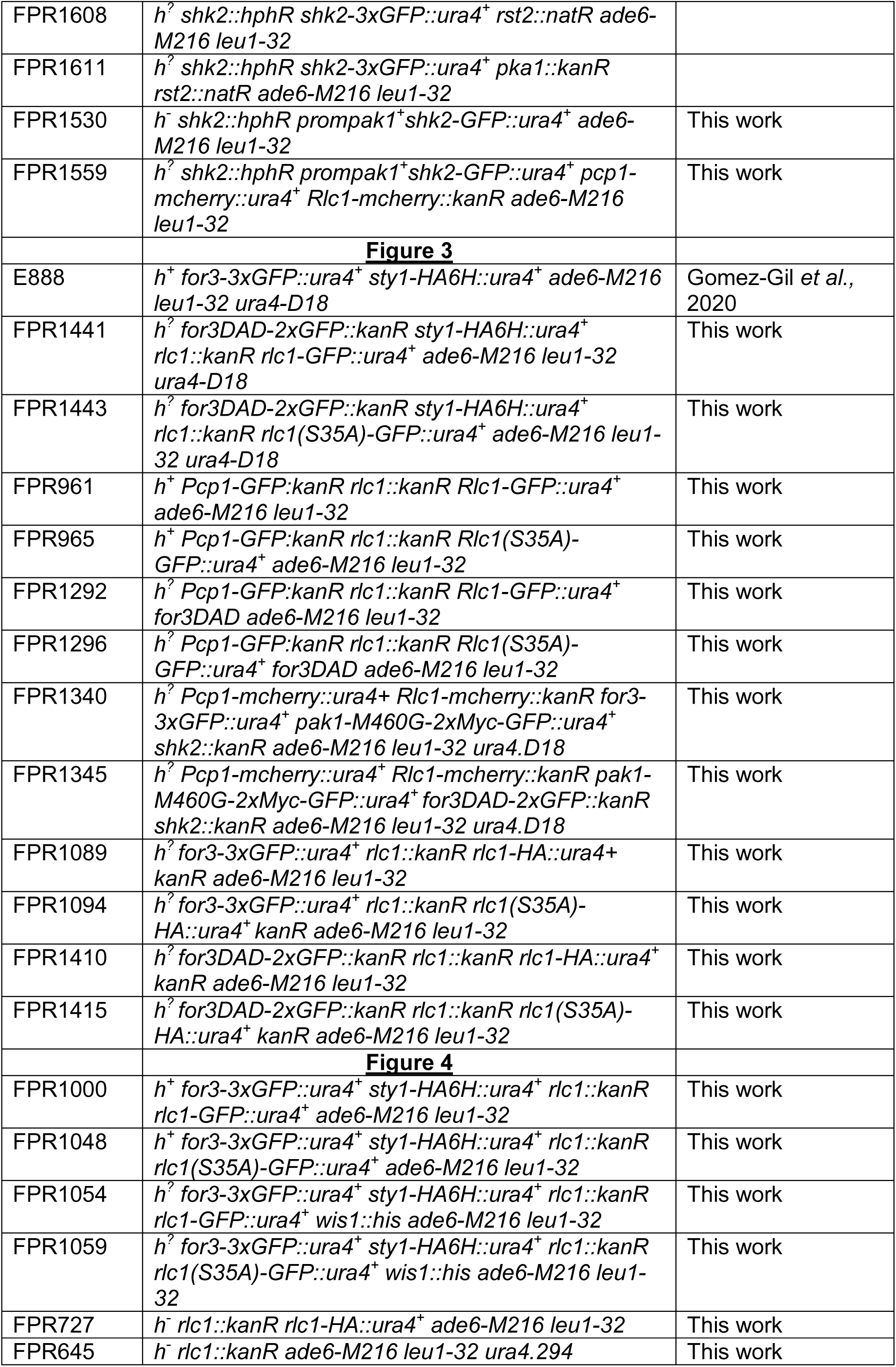
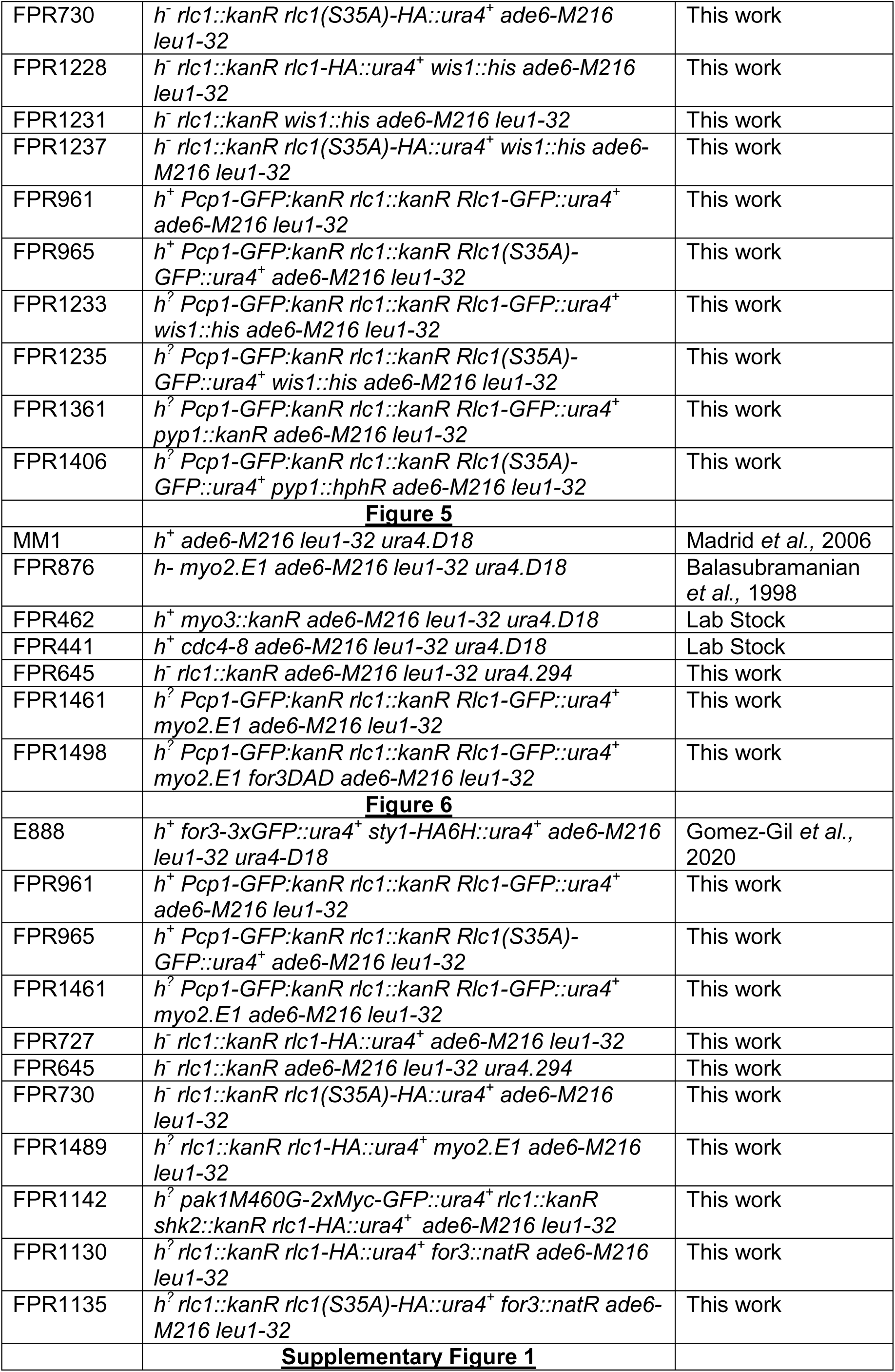
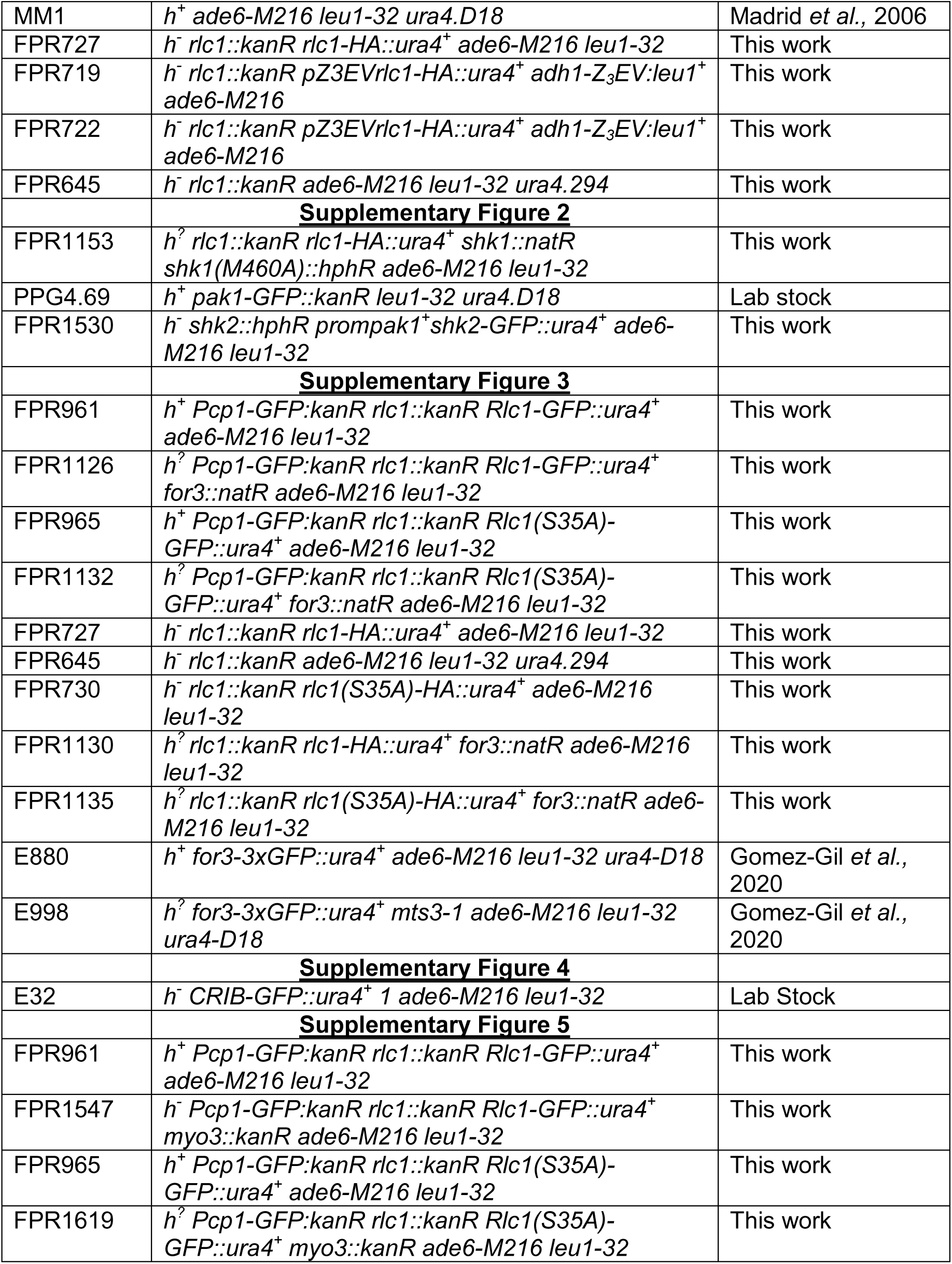
*S. pombe* strains used in this study.

**Table S2.**
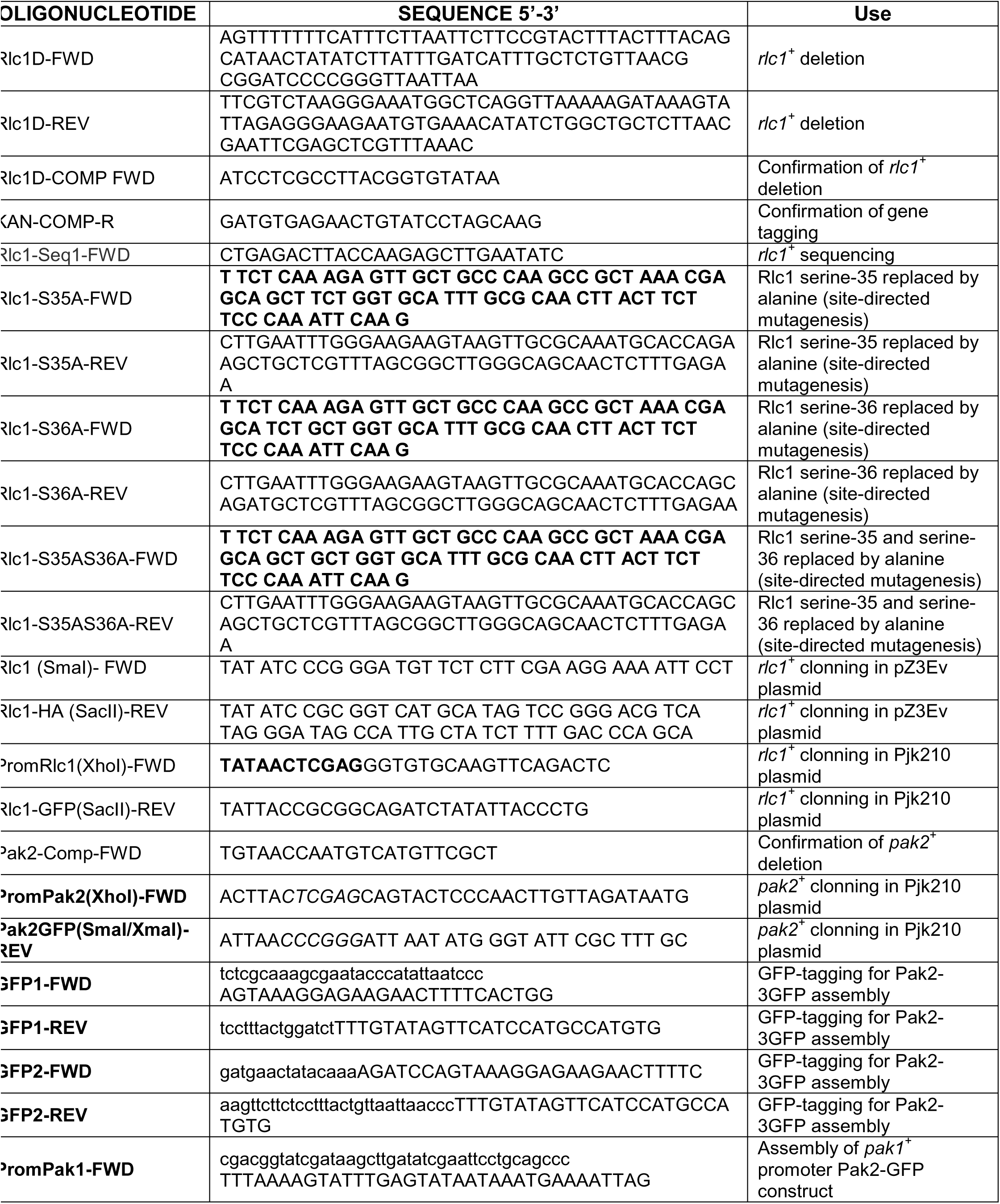
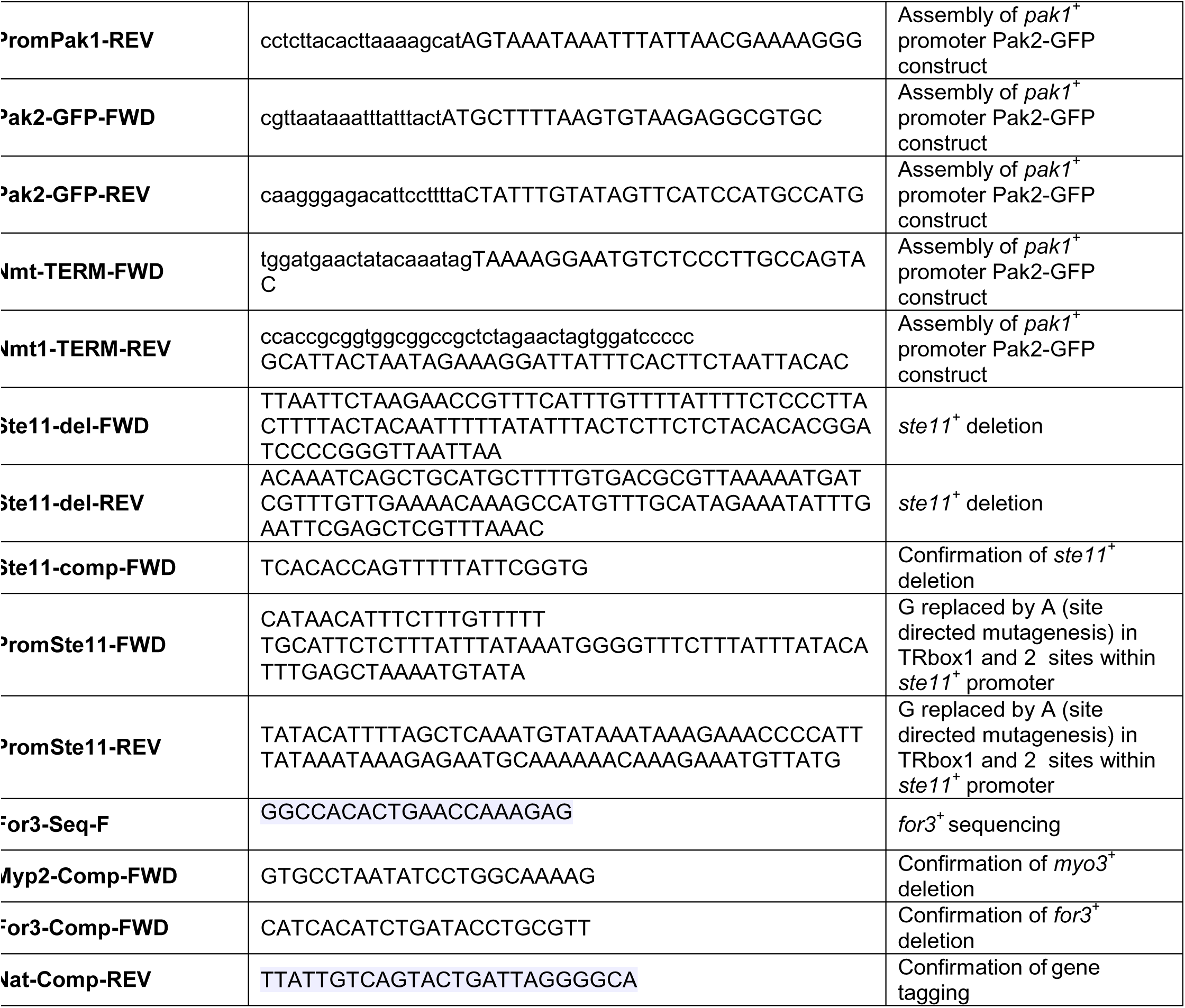
Oligonucleotides and DNA fragments used in this study.

